# Surface proteomics reveals arginine metabolism as a vulnerability in high grade serous ovarian cancer

**DOI:** 10.1101/2025.09.02.673726

**Authors:** Meinusha Govindarajan, Matthew Waas, Salvador Meija-Guerrero, Brian Lin, Tianyi Ling, Ximing Li, Vladimir Ignatchenko, Anastasia N. Tikhonova, Courtney L. Jones, Laurie Ailles, Thomas Kislinger

## Abstract

**Background:** The significant lethality of high-grade serous ovarian cancer (HGSC) is driven by the lack of long-term efficacy of current treatments, underscoring the need for developing additional therapeutics. Cell surface proteins represent attractive therapeutic targets yet remain underexplored in high-grade serous ovarian cancer. Here, we employed cell surface *N*-glycoproteomics to elucidate the cell surface proteomes of HGSC cells alongside normal epithelial and cancer-associated stromal cells, uncovering new opportunities for therapeutic intervention.

**Results:** Integration of cell surface *N*-glycoproteomics and functional screening revealed multiple minimally characterized, HGSC-enriched surface proteins that are critical for HGSC proliferation – most notably, SLC7A1. Multi-omic and functional characterization indicated that SLC7A1 is necessary for HGSC migration, protein synthesis and mitochondrial functions likely linked to its role as an arginine transporter. Finally, we demonstrate that elevated surface expression of SLC7A1 in HGSC reflects dysregulated arginine metabolism, pointing to a putative therapeutic vulnerability.

**Conclusions:** Our work identifies SLC7A1 and arginine metabolism as a previously unrecognized molecular vulnerability in HGSC and provides a framework to guide future therapeutic development.

## BACKGROUND

High-grade serous ovarian cancer (HGSC) is the most lethal gynecological malignancy in Canada, the United States, and Europe (1–3). The current standard of care involves debulking surgery followed by platinum-based combination chemotherapy (4). While initial remission is often achieved, most patients experience a relapsing course of disease with escalating chemoresistance, ultimately leading to death (4). Recent targeted therapies, such as a FOLR1-directed antibody-drug conjugate (ADC) (5) and multiple PARP inhibitors (6,7), have shown promise for select patient subsets, yet the five-year overall survival for HGSC patients remains dismal at approximately 30% (8). As HGSC is still terminal for most patients, the development of additional targeted therapies is vital to help control disease and prolong survival.

Cell surface proteins are a valuable class of proteins comprising approximately 60% of all FDA approved protein drug targets (9). Their imperative roles as gatekeepers between intra- and extracellular signaling, coupled with their accessible outer surface localization establishes them as attractive candidates for therapeutic targeting (10). Characterization of cell surface proteins associated with malignancy can thus enable the identification of targets exploitable by novel therapies. However, defining the surface-exposed proteome (*i.e.,* the surfaceome) is challenging as widely used high-throughput methods (*e.g.,* transcriptomics, global proteomics, antibody screening) are not optimally suited for accurately profiling cell surface proteins (10,11). These limitations arise from several challenges including a sub-optimal correlation between RNA expression and surface protein abundance, the under-representation of surface proteins in global proteomics experiments due to their low abundance and biophysical properties and the limited availability of validated antibodies (10,12). As a result, comprehensive surfaceome resources remain absent for many disease contexts, including HGSC, limiting the discovery of new therapeutic modalities.

Here, we utilized Cell Surface Capture (CSC) (13) – a highly specific surface *N*-glycoproteomic method – to elucidate the *in vitro* HGSC surfaceome and compare it to that of normal epithelial and cancer-associated stromal cells. Functional screening of HGSC enriched surface proteins uncovered multiple novel surface proteins critical for HGSC proliferation, including arginine transporter SLC7A1. Detailed proteomic and phenotypic interrogation of SLC7A1 knockdown in HGSC revealed impaired migration, protein synthesis, and cell survival. Finally, we show that arginine metabolism is dysregulated in HGSC cells compared to their normal counterparts, underscoring a putative actionable vulnerability for HGSC.

## RESULTS

### Characterization of the *in vitro* HGSC surface proteome

Leveraging the high prevalence of *N*-glycoproteins on the extracellular surface (14,15), we employed CSC – an established method for enriching surface exposed *N*-glycoproteins (13) – on a panel of nine models to empirically define the cell surface proteome of HGSC and related cell types (**Figure 1A**). Specifically, we profiled four commercially available HGSC epithelial cell lines (OVCAR8, Kuramochi, PEO4 and ES2) selected for their close molecular resemblance to HGSC patient tumors (16,17), two normal fallopian tube epithelial (FTE) cell lines (FT237 and FT194) that represent the cell of origin for most HGSC as non-cancerous controls (18,19), and three HGSC patient-derived cancer-associated fibroblast (CAF) populations (CAF3028, CAF438 and CAF40879) as stromal controls (20). Surface proteome profiling was highly reproducible as evidenced by strong correlation between cell line replicates (Median = 0.87, **Figure S1A, B**). In total, we detected 855 extracellularly exposed *N*-glycoproteins **(Additional File: Table S1).** 79% of detected *N*-glycoproteins were predicted surface membrane proteins and 86% contained a predicted signal peptide (14) (**Figure 1B**). Moreover, 93% of detected *N*-glycopeptides mapped to the predicted extracellular domain of their respective proteins, highlighting the specificity of CSC for profiling plasma membrane and secreted proteins.

**Figure 1:**
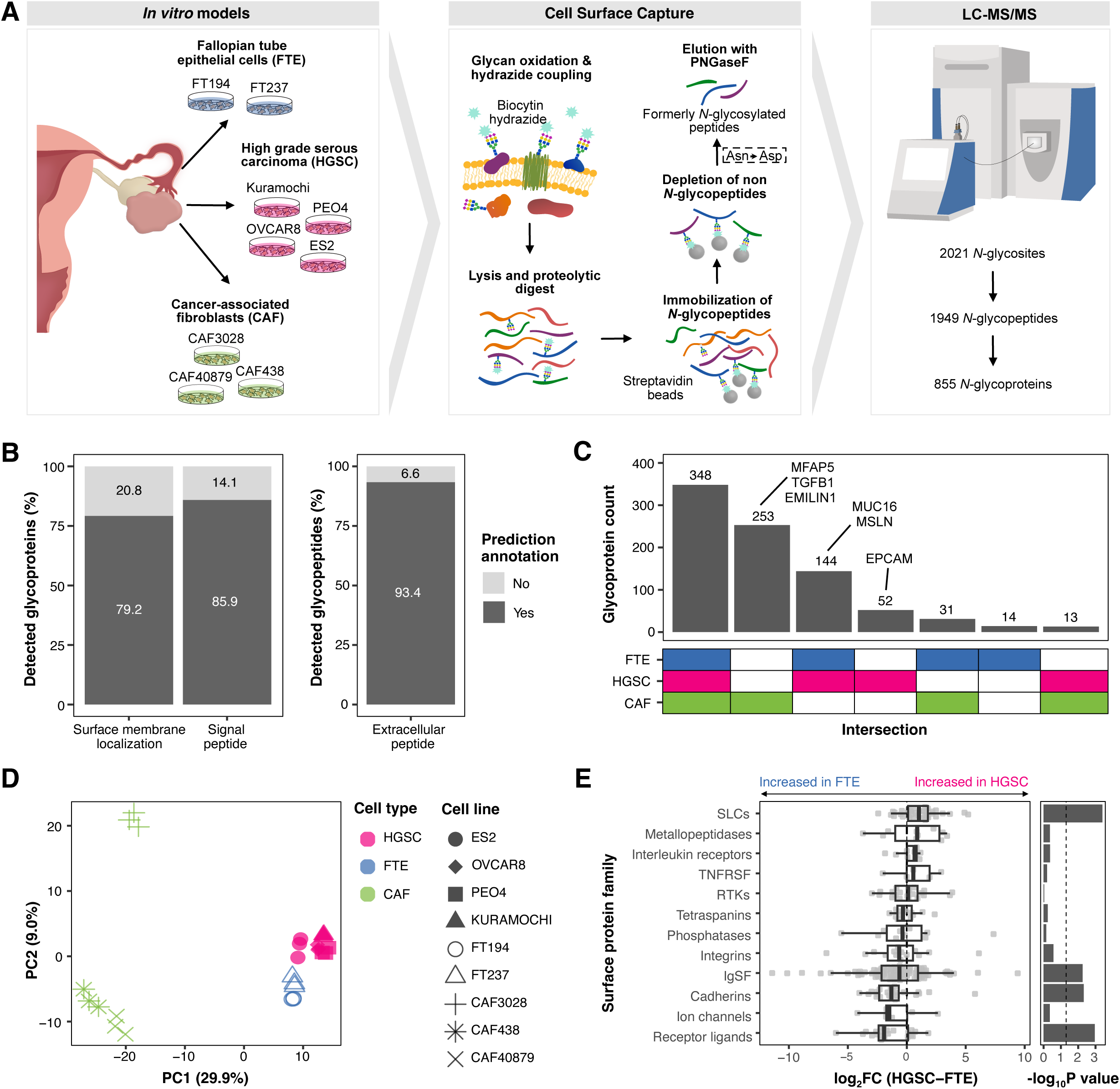
Surface proteomic characterization of HGSC. **A)** Overview of cell surface capture (CSC) *N*-glycoproteomic approach used to profile the surfaceome of HGSC cells (n=4), FTE cells (n=2) and CAFs (n=3). Three processing replicates per model. **B)** Percentage of total detected *N*-glycoproteins (left) and *N*-glycopeptides (right) with predicted surface and secreted protein annotations (14). **C)** Upset plot indicating uniquely detected and shared proteins between the surfaceomes of HGSC, FTE and CAF cell types. Known cell type associated surface proteins are annotated. **D)** Principal component analysis of CSC *N*-glycoproteomic data. Shape indicates cell line and color denotes cell type. Each point represents a cell line processing replicate (n=3). **E)** Differential surface abundances between HGSC and FTE cells categorized by HGNC surface protein families. Boxplots of log_2_fold changes (left) and -log_10_p-values determined by a one-sample Mann-Whitney U-test (right) for each protein family are visualized.

The highest number of extracellularly exposed *N*-glycoproteins were detected in CAFs, followed by FTE cells and lowest in HGSC cells (**Figure S1C**). As CAFs are defined by a highly secretory phenotype (21) and FTE cells are of a secretory epithelial lineage (18), we hypothesized that these differences may be reflective of variations in secretory capacity. Indeed, analyzing our dataset by subcellular localization (*i.e.,* plasma membrane *vs*. secreted) revealed notable cell type differences in the detection of secreted proteins (∼2X difference between cell types) but not plasma membrane proteins (∼1.2X difference between cell types) **(Figure S1D**). Furthermore, cell type explained substantially more variance in secreted protein detection (ω^2^ = 0.89) than plasma membrane protein detection (ω^2^ = 0.22), consistent with differences in secretory capacity across cell types contributing to the observed variation in total detection depth. Analysis of cell type unique and shared surface proteins revealed a greater overlap between HGSC and FTE cells than with CAFs (**Figure 1C**). This is consistent with their shared epithelial lineage, in contrast to the mesenchymal origin of CAFs. Notable HGSC-associated surface proteins, MUC16 and MSLN, were amongst the 144 proteins shared between HGSC and FTE cells. Despite the high degree of overlap between HGSC and FTE cells, principal component analysis illuminated quantitative differences between the surfaceomes of malignant and non-cancerous cells (**Figure 1D**). HGSC cells had an increased abundance of solute carriers and a decreased abundance of receptor ligands, cadherins, and members of the immunoglobulin superfamily compared to FTE cells (**Figure 1E**). There was also a decreased abundance of integrins, tetraspanins and receptor ligands on the surface of HGSC cells compared to CAFs (**Figure S1E**). Collectively, our analysis reveals distinct cell surface repertoires across *in vitro* HGSC, FTE, and CAF populations, with malignant cells characterized by solute carrier enrichment and loss of adhesion molecules, highlighting unique therapeutic opportunities in ovarian cancer.

### Functional screening reveals surface proteins critical for HGSC proliferation and survival

With the goal of identifying HGSC enriched surface proteins, we integrated our *in vitro* surface proteomics data with patient global proteomics data and devised a target prioritization strategy (**Figure 2A**). First, we filtered our data to retain 274 high-confidence HGSC surface proteins (*i.e.,* proteins that were robustly detected on the surface of ≥ 3 HGSC cell lines and in ≥1 HGSC patient tumor profiled by the Clinical Proteomic Tumor Analysis Consortium [CPTAC] (22,23)). Next, we developed a target score (see **Methods**) which prioritized proteins based on three criteria: 1) increased surface abundance in HGSC cells compared to FTE cells and CAFs; 2) frequent detection in the proteomes of HGSC patient tumors (22,23); and 3) limited detection in normal tissue found throughout the human body (9,24–26). Ranking the 274 high-confidence surface proteins based on our target score revealed FOLR1 – the only FDA approved ADC target currently available for HGSC (5) – amongst the highest ranked proteins, confirming our prioritization scheme as an effective method for identifying clinically relevant HGSC proteins (**Figure 2B**). While some of the highly ranked proteins had already been extensively characterized in the context of ovarian cancer (*e.g.,* SLC2A1 (27–32) and TFRC (33–37)), we noticed there were sparse reported associations with HGSC for several other top ranked proteins, suggesting the elucidation of potentially novel HGSC surface markers (**Additional File: Table S2**).

**Figure 2:**
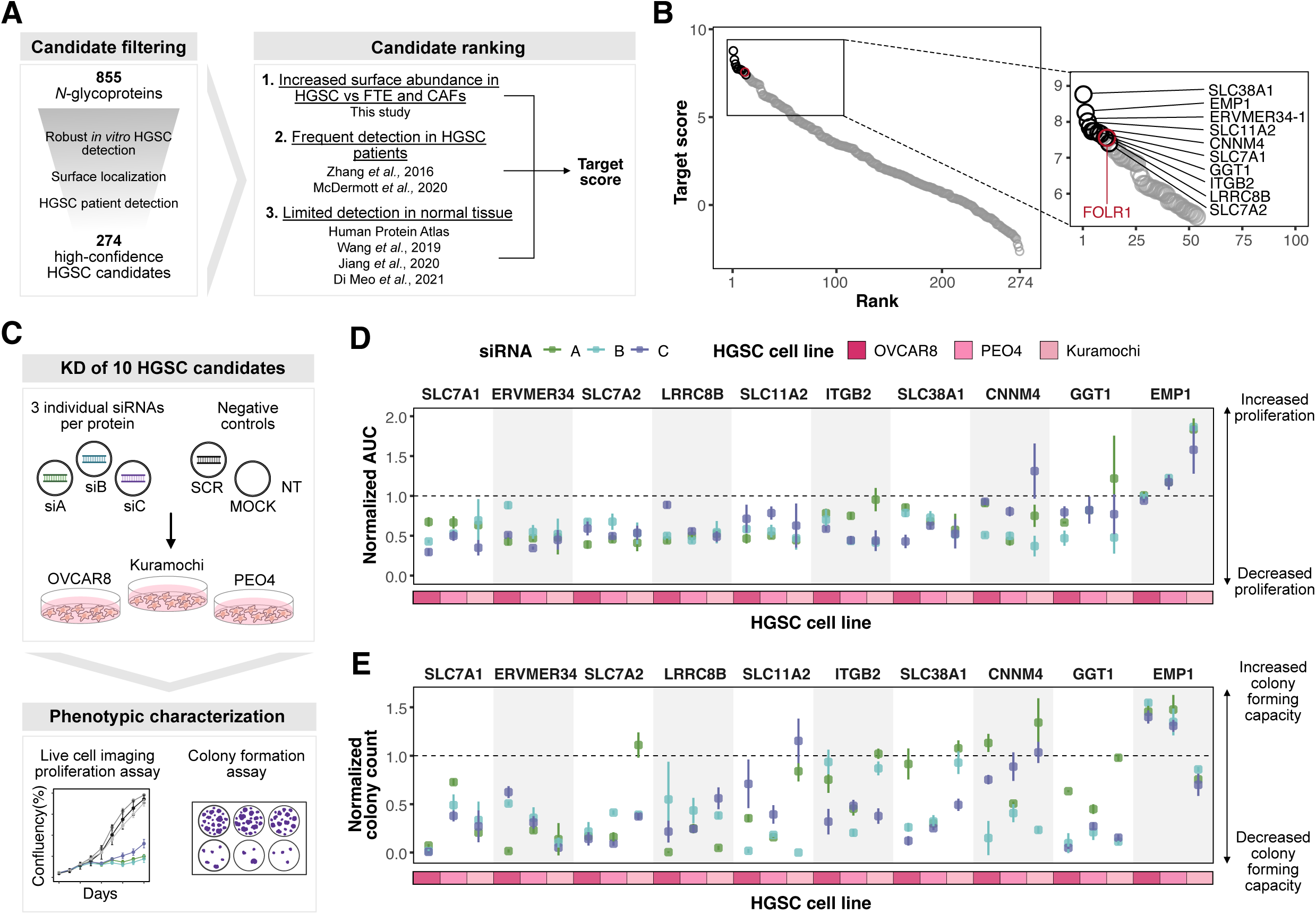
Functional evaluation of HGSC surface proteins reveals actionable vulnerabilities. **A)** Strategy to filter and prioritize HGSC enriched surface candidates. **B)** 274 HGSC enriched surface candidates ranked based on the target score devised in **(A)**. FOLR1 and ten HGSC surface proteins selected for functional evaluation are annotated in the zoom-in panel. **C)** Schematic of siRNA screen used to functionally evaluate ten HGSC surface proteins. **D)** Area under the proliferation curves (AUC) normalized to scramble for HGSC cells treated with one of three siRNAs per target. Data are represented as mean (n=3) ± SD. **E)** Colony counts normalized to scramble for HGSC cells treated with one of three siRNAs per target in three cell lines. Data are represented as mean (n=3) ± SD.

Consequently, we selected 10 highly ranked HGSC enriched surface proteins with relatively scarce or preliminary literature pertaining to HGSC (SLC38A1, EMP1, ERVMER34-1, SLC11A2, CNNM4, SLC7A1, GGT1, ITGB2, LRRC8B, SLC7A2) for functional screening. Notably, 6/10 of the HGSC target candidates prioritized for functional interrogation were undetected in previous global proteomics analyses of these models (38), underscoring the unique discovery potential of cell surface proteomics approaches. For each target, we evaluated the effect of three independent siRNAs on *in vitro* cell growth and survival using live-cell imaging and colony forming assays (**Figure 2C**). Knockdown of 8/10 selected HGSC enriched surface proteins had deleterious effects on proliferation, as measured by live-cell imaging, compared to a scramble siRNA (**Figure 2D**). For four candidates, siRNA transfection also consistently impaired colony forming capacity across all three HGSC cell lines (**Figure 2E**), indicating that these proteins are important for cell survival. Altogether, we have identified four minimally characterized, HGSC enriched surface proteins (SLC7A1, ERVMER34-1, SLC7A2 and LRRC8B) that are robustly critical for HGSC proliferation and survival, providing new insights into HGSC dependencies.

### SLC7A1 is a cancer enriched surface protein essential for HGSC growth

High-affinity cationic amino acid transporter 1 (SLC7A1) was selected for additional interrogation due its intriguing cancer enriched expression profile and its strong essentiality across multiple HGSC cell lines. Our *in vitro* data demonstrated that SLC7A1 is highly abundant on the surface of HGSC cells compared to normal FTE cells and CAFs (3.2 - 4.9X greater, **Figure 3A**). This cancer enriched expression profile was corroborated in a patient proteomics dataset (comparing bulk HGSC tumors and fallopian tubes) (23) (**Figure 3B**) and a spatial transcriptomic patient dataset (comparing the epithelial components of HGSC tumors to HGSC precursor lesions and normal fallopian tubes) (39) (**Figure S2A**). As a small proportion of HGSC tumors are thought to arise from the ovarian surface epithelium (40,41), we also queried a patient proteomics dataset comparing ovarian tumors to benign ovarian masses and normal ovaries (42) and observed similar cancer enriched protein abundance of SLC7A1 (**Figure S2B**). Evaluation of an in-house HGSC patient tumor scRNA-seq dataset verified higher expression of *SLC7A1* in HGSC epithelial cells compared to other cells in the tumor microenvironment, including fibroblasts (**Figure S2C**). Interestingly, high expression of SLC7A1 was associated with poorer overall survival in the TCGA RNA-seq cohort of HGSC patients (43) using both univariate (HR = 1.36, P = 0.038, **Figure 3C**) and multivariate Cox regression adjusting for age and FIGO stage (HR = 1.37, P = 0.036, **Figure S2D)**. Regarding systemic normal tissue abundance, SLC7A1 was only detected in 8/32 normal tissues profiled by mass spectrometry – among which, the salivary gland was the only tissue with a frequency of detection above 10% (24) (**Figure 3D**). Analysis of an independent proteomics dataset by Human Protein Atlas (44) and a RNA-seq dataset generated by GTEx (45) also confirmed restricted normal tissue expression of *SLC7A1* (**Figures S2E & F**). Our functional screen (**Figure 2C**) revealed that knockdown of SLC7A1 with three independent siRNAs (**Figure S3A**) impaired *in vitro* HGSC cell proliferation (**Figures 3E & S3B**) and colony forming capacity (**Figures 3F & S3C**). The observed phenotypes were consistent using all three siRNAs in three HGSC cell lines. To further evaluate SLC7A1 essentiality using a stable loss-of-function approach, we generated doxycycline (dox)-inducible shRNA-containing OVCAR8 cells targeting SLC7A1 with two independent hairpins **(Figure S3D)**. Consistent with our transient siRNA data, shRNA-mediated depletion of SLC7A1 markedly impaired HGSC proliferation and colony formation capacity compared to a GFP-targeting shRNA control **(Figures S3E-G)**. To assess the consequences of SLC7A1 depletion *in vivo*, dox-inducible shRNA OVCAR8 cells were subcutaneously engrafted into immunocompromised mice, with dox administered in drinking water 10 days post-inoculation to enable *in vivo* knockdown. Notably, induction of SLC7A1 shRNAs arrested tumor growth compared to GFP shRNA controls **(Figures 3G & H)**. Together, these data identify SLC7A1 as a cancer-enriched surface protein in HGSC warranting mechanistic investigation into its contribution to tumor growth.

**Figure 3:**
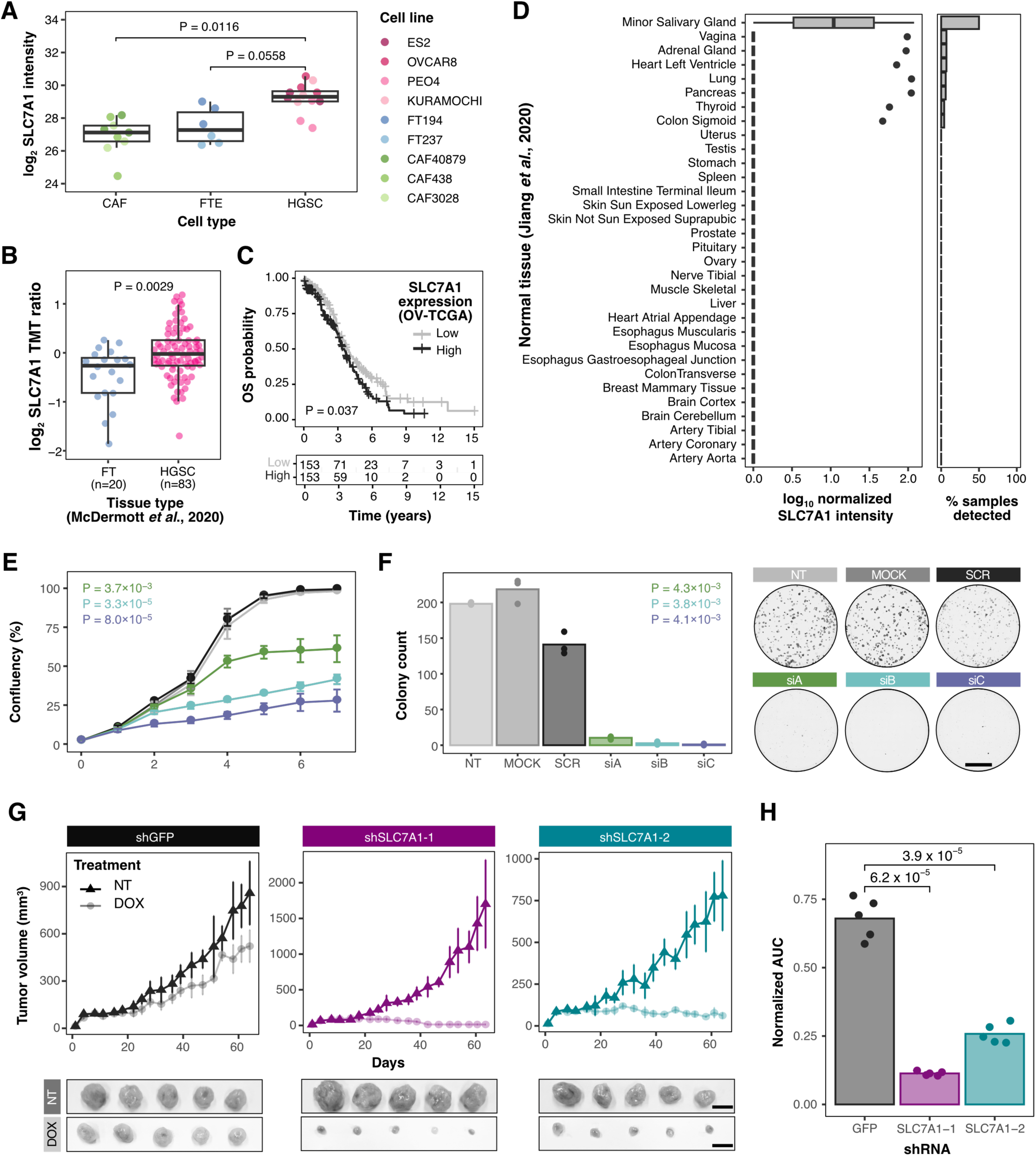
SLC7A1 is a cancer enriched surface protein essential for HGSC growth. **A)** SLC7A1 protein abundance on the cell surface of HGSC, FTE and CAF cells as determined by mass spectrometry. P-values determined using a linear-mixed effects model (intensity ∼ cell type + (1|cell line)). Three replicates per cell line. **B)** SLC7A1 protein abundance in HGSC and normal fallopian tube (FT) patient tissue profiled by McDermott *et al.,* 2020 (23). P-value was determined using a two-sample Mann-Whitney U-test. **C)** Univariate overall survival based on median dichotomized SLC7A1 expression in HGSC tumors profiled by TCGA (43). Statistical significance was calculated with a log-rank test. **D)** SLC7A1 protein abundance (left) and detection frequency (right) in normal tissues profiled by Jiang *et al.,* 2020 (24). **E)** Proliferation of OVCAR8 cells treated with three independent siRNAs against SLC7A1. Data are represented as mean ± SD (n=3). Statistical significance was determined by comparing the areas under the proliferation curves for each siRNA against SCR with a Student’s t-test. **F)** Quantification (left) and representative images of colony formation assays (right) following transient transfection with SLC7A1 targeting siRNAs. Means and individual data points (n=3) are plotted. P-values were calculated using a Student’s t-test against SCR. Scale bar = 10,00 µM. **G)** *In vivo* growth curves (top) and endpoint images (bottom) for subcutaneous OVCAR8 xenografts with doxycycline (dox)-inducible shRNAs. shGFP was used as a non-targeting negative control condition. Scale bar = 1 cm. **H)** Areas under the proliferation curve (AUC) relative to respective non-treated (NT) cells in each condition from **(G)** (means and individual datapoints, n=5). P-values were determined using a Student’s t-test.

### Knockdown of SLC7A1 dysregulates several vital cellular processes in HGSC

To explore the function of SLC7A1 in HGSC models, we performed whole cell lysate proteomics on OVCAR8 cells treated with one of three siRNAs against SLC7A1 or a scrambled siRNA (**Figure 4A**). LC-MS/MS analysis confirmed strong protein knockdown (>90%) of SLC7A1 with all three siRNAs compared to scramble 72h post transfection (**Figures 4B & S4A**). Proteomic pathways that were consistently dysregulated across all three siRNAs related to cell growth and migration, protein regulation, DNA synthesis, mitochondrial functions, and cell death (**Figure 4C, Additional File: Table S3**). 3D spheroid and scratch-wound migration assays confirmed that SLC7A1 knockdown impaired OVCAR8 spheroid growth (**Figure 4D**) and migration (**Figure 4E**), respectively. Measurement of puromycin incorporation into newly translated proteins indicated decreased protein synthesis as a function of SLC7A1 downregulation **(Figure S4B)**. Evaluation of 5-ethynyl-2’-deoxyuridine (EdU) incorporation into newly synthesized DNA revealed reduced DNA synthesis following SLC7A1 knockdown (**Figure S4C**). Incubation of OVCAR8 cells with MitoSOX and tetramethylrhodamine ethyl ester (TMRE) demonstrated increased mitochondrial ROS (**Figure S4D**) and membrane polarization (**Figure S4E**), respectively, corroborating dysregulated mitochondrial function following SLC7A1 depletion. Finally, assessment of Annexin V positivity and caspase activation – markers of apoptosis – verified caspase mediated OVCAR8 cell death as a consequence of SLC7A1 downregulation (**Figures 4F**, **S4F & S4G**). Overall, our proteomic and functional characterization uncovered that SLC7A1 depletion alters cell growth, migration, mitochondrial function, DNA and protein synthesis, culminating in HGSC cell death.

**Figure 4:**
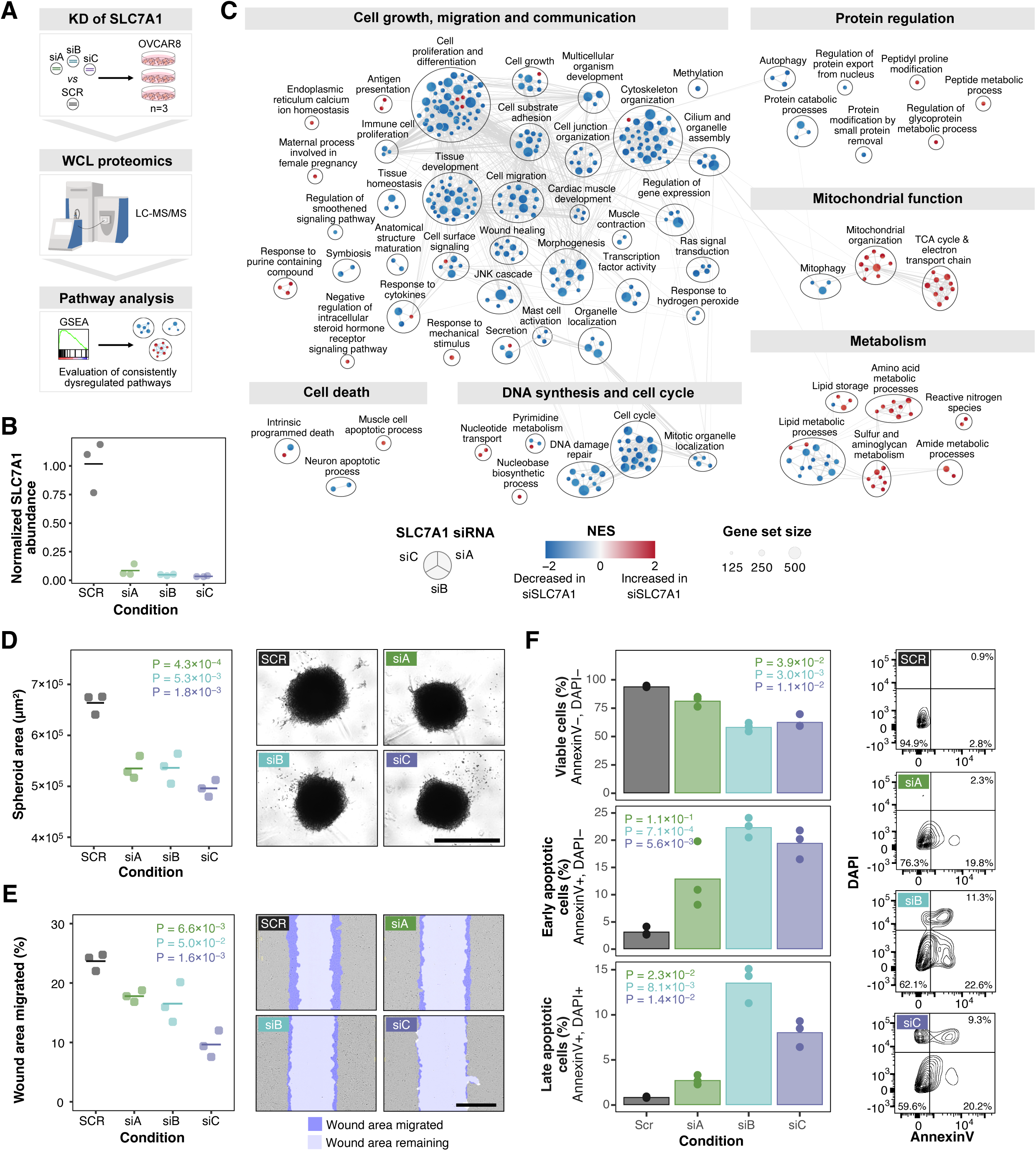
SLC7A1 knockdown impairs several cellular processes in HGSC. **A)** Schematic of whole-cell lysate (WCL) proteomics analysis performed on OVCAR8 cells treated with SLC7A1 targeting siRNAs for 72 h. Three processing replicates per condition. **B)** SLC7A1 protein abundance in OVCAR8 cells 72h following siRNA treatment, normalized to cells treated with a Scramble (SCR) siRNA. **C)** Enrichment map showing GO:Biological pathways consistently dysregulated across all three independent SLC7A1 targeting siRNAs in OVCAR8 cells. **D)** Quantification (left) and representative images (right) of OVCAR8 spheroids 10 days post transfection with SLC7A1 or SCR siRNAs. Student’s t-test P-values against SCR are reported. Scale bar = 1000 µM. **E)** Quantification (left) and representative images (right) of scratch-wound migration assays performed in OVCAR8 cells 24h post treatment with SLC7A1 or SCR siRNAs. Student’s t-test P-values against SCR are reported. Scale bar = 600 µM. **F)** Quantification of percentage viable (top), early apoptotic (middle) and late apoptotic (bottom) OVCAR8 cells 96h post transfection with SLC7A1 or SCR siRNAs. Representative flow cytometry contour plots for each condition are shown on the left. Student’s t-test P-values against SCR are reported. (**B, D-F**) Means and individual data points (n=3) are plotted.

### Arginine metabolism is a molecular vulnerability in HGSC

As SLC7A1 is canonically known to be an arginine (Arg) transporter (46), we postulated that our observed phenotypes may be linked to reduced Arg availability in the cell. To this end, we pulse labelled OVCAR8 cells with ^13^C_6_-Arg to evaluate Arg influx following 48 h of SLC7A1 depletion (**Figure 5A**). LC-MS/MS metabolomic analysis revealed reduced Arg uptake in OVCAR8 cells treated with SLC7A1 siRNAs compared to a scrambled siRNA (**Figures 5A & S4H**). Immunoblotting also indicated that SLC7A1 knockdown correlated with a time-dependent decrease in mTORC1 signaling (**Figure 5B**), a pathway that requires Arg for activation (47–49). Notably, depletion of Arg from cell culture media (*i.e*., media containing 0% Arg) impaired OVCAR8 proliferation (**Figure 5C**) similar to that of SLC7A1 knockdown (**Figure 3E**). Live-cell imaging also demonstrated a dose-response relationship between the level of Arg in cell culture media and OVCAR8 proliferation. This relationship was most prominently observed at Arg media concentrations lower than 115 µM (*i.e.,* 10% of Arg found in complete media [CM]) which is roughly equivalent to the basal amount of Arg found in human blood circulation (50,51), highlighting that OVCAR8 cells were sensitive to reductions in extracellular Arg within the physiologic range. Overall, these findings suggest that the proliferative defects following SLC7A1 loss was associated with impaired Arg availability, prompting us to examine the broader role of Arg metabolism in sustaining HGSC growth.

**Figure 5:**
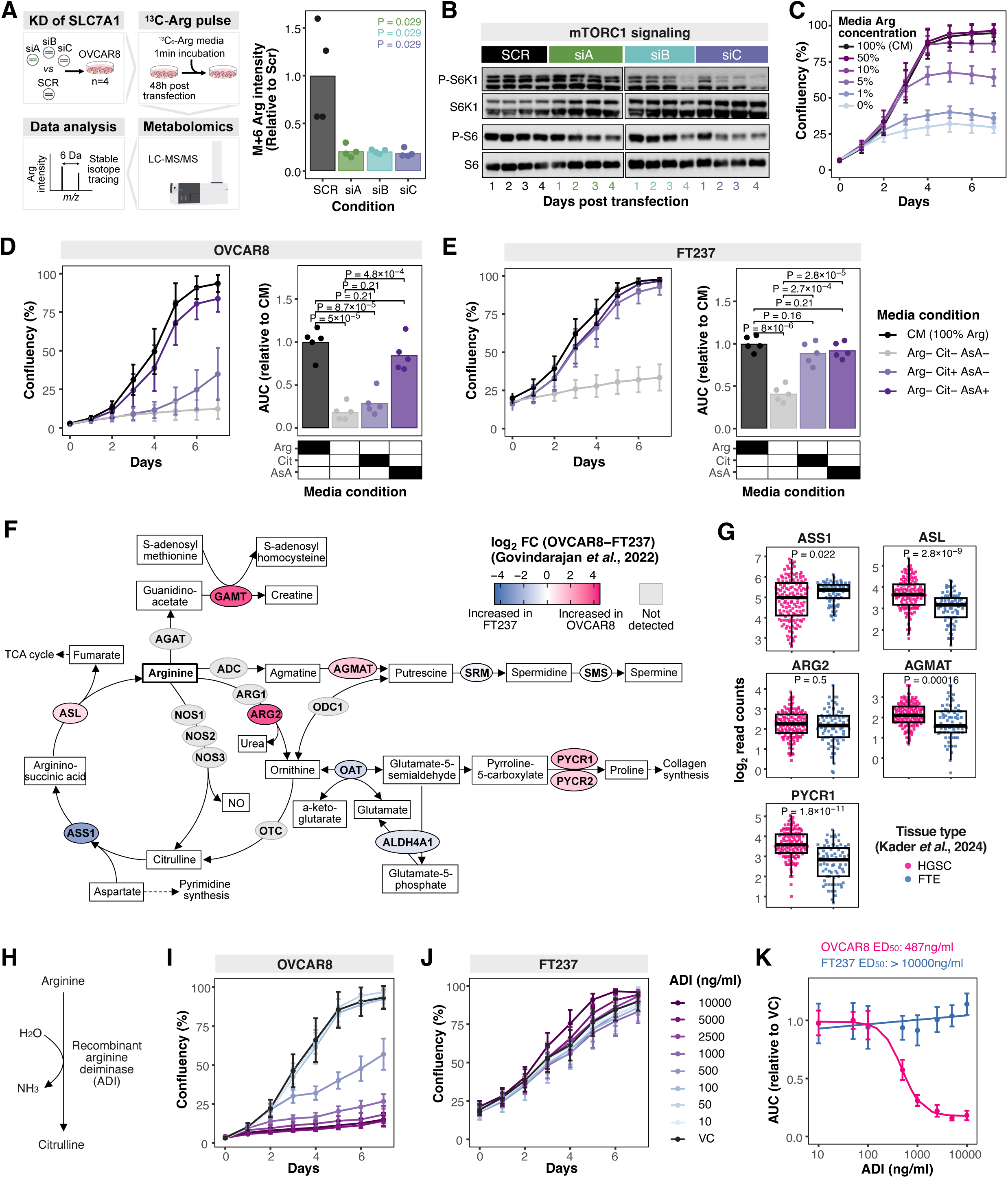
Arginine metabolism is a molecular vulnerability in HGSC. **A)** Stable-isotope tracing metabolomics performed on OVCAR8 cells treated with SLC7A1 or Scramble (SCR) siRNAs. Experimental schematic (left) and intensity of M+6 (*i.e.,* heavy labelled) Arg relative to SCR (right) are visualized. Means and individual datapoints (n=4) are plotted. P-values were calculated using a two-sample Mann-Whitney U-test against SCR. **B)** Time-course immunoblotting of mTORC1 signaling in OVCAR8 cells following transfection with SLC7A1 or SCR siRNAs. **C)** Proliferation of OVCAR8 cells grown in RPMI 1640 media containing varying concentrations of Arg. Means and SDs (n=5) are plotted. Complete media (CM) refers to media containing the amount of Arg regularly found in RPMI 1640 (*i.e.,* 1.15 mM Arg or 100% Arg). **D)** Proliferation curves (left) comprised of means and SDs (n=5) for OVCAR8 cells grown in different media compositions. Areas under the proliferation curves (AUC) relative to CM are plotted (means and individual datapoints) on the right. P-values were determined using a Student’s t-test. **E)** Proliferation curves (left) comprised of means and SDs (n=5) for FT237 cells grown in differing media compositions. AUCs relative to CM are plotted (means and individual datapoints) on the right. P-values were determined using a Student’s t-test. **F)** Overview of arginine metabolism. Metabolism enzymes are shaded based on proteomic log_2_ fold changes between FT237 and OVCAR8 cells profiled in Govindarajan *et al.,* 2022 (38). Light grey indicates not detected. **G)** Expression of ASS1, ASL, ARG2, AGMAT and PYCR1 in HGSC and FTE patient tissue profiled by Kader *et al.,* 2024 (39). Each point represents an independent epithelial section. P-values were calculated using a two-sample Mann-Whitney U-test. **H)** Schematic of arginine depletion mediated by arginine deiminase (ADI). **I)** Proliferation curves of OVCAR8 cells treated with varying concentrations of ADI. Means and SDs (n=5) are plotted. **J)** Proliferation curves of FT237 cells treated with varying concentrations of ADI. Means and SDs (n=5) are plotted. **K)** ADI dose-response curves for OVCAR8 and FT237 cells fitted based on the AUCs derived from (**I&J**).

Arg is theoretically a non-essential amino acid because human cells contain the machinery to synthesize Arg from other metabolites (*i.e.,* citrulline [Cit] and argininosuccinic acid [AsA]) (52). However, under certain conditions, such as times of rapid growth or stress, cells cannot produce enough Arg to sustain cellular activities and instead, rely on transporting Arg into the cell from extracellular sources (52). As a result, Arg can be considered a “conditionally essential” or “semi essential” amino acid in practice (52). To test Arg essentiality in HGSC models, we evaluated whether the presence of Cit or AsA (*i.e.,* Arg precursor metabolites) in cell culture media could restore OVCAR8 proliferation in the absence of Arg. While the addition of AsA mostly rescued the negative effects of Arg starvation, supplementation with Cit only marginally improved cell proliferation and did not restore proliferation rates to that of OVCAR8 cells grown in complete media (CM, *i.e.,* media containing 100% Arg) (**Figure 5D**). We next assessed whether this phenomenon was cancer-specific and found that unlike OVCAR8 cells, the presence of Cit or AsA in media was sufficient to completely maintain the proliferative capacity of FT237 cells in the absence of Arg (**Figure 5E**). Evaluation of our previously published whole cell lysate proteomics dataset on OVCAR8 and FT237 cells (38) provided insight into the differential essentiality of Arg between these two cell types, as several enzymes involved in Arg synthesis and utilization were differentially abundant (**Figure 5F**). Expression differences between HGSC and FTE of select enzymes were corroborated in a spatial transcriptomics patient dataset (39) (**Figure 5G**). Most notably, argininosuccinate synthase (ASS1) – the enzyme which metabolizes Cit – was found to be downregulated in HGSC compared to normal FTE. Together, these data suggest that Arg is semi essential for OVCAR8 cells and non-essential for FT237 cells and point towards altered Arg metabolism between HGSC and FTE cells.

Therapeutically targeting Arg metabolism with Arg-depleting enzymes has emerged as a promising strategy in several cancers (53). Specifically, formulations of arginine deiminase (ADI), a recombinant bacterial enzyme that converts Arg into Cit (**Figure 5H**), are currently under clinical trial for treating hepatocellular carcinoma, mesothelioma and melanoma (53). Provided that Cit demonstrated a differential ability to sustain the proliferative capacity of OVCAR8 and FT237 cells following Arg depletion (**Figures 5D&E**), we hypothesized that HGSC and FTE cells may exhibit distinct sensitivity to ADI. Indeed, treatment with ADI resulted in a dose-dependent decrease in OVCAR8 proliferation and had no notable effects on FT237 proliferation at all tested concentrations, spanning four orders of magnitude (**Figures 5I-K**). These data demonstrate that arginine metabolism may represent an actionable molecular vulnerability for HGSC.

## DISCUSSION

HGSC remains the most aggressive and lethal form of epithelial ovarian cancer (EOC) largely due to late-stage diagnosis and limited long-term efficacy of current therapeutic strategies (4). Only in the last decade has the FTE been widely accepted as the primary origin of HGSC (54), leaving a significant gap in molecular comparisons between HGSC and its normal counterpart. Moreover, the cell surface – a clinically relevant, context specific subcellular compartment – of HGSC remains poorly defined due to challenges associated with comprehensively profiling the cell surface proteome. Here, we utilized CSC, a highly specific *N*-glycoproteomic surface profiling technique (13), to elucidate the first surfaceome maps of HGSC, non-cancerous FTE and HGSC CAFs. Coupling our surface proteomics data with functional screening uncovered four understudied, cancer enriched surface proteins critical for HGSC growth and survival (SLC7A1, ERVMER34-1, SLC7A2 and LRRC8B). SLC7A1 and SLC7A2 are solute carrier transporters of cationic amino acids (46). ERVMER34-1 is an endogenous retroviral envelope protein whose function in the human proteome remains largely uncharacterized (55). LRRC8B is a subunit of volume-regulated anion channels involved in cell volume regulation and osmotic stress response (56). As cancer enriched surface proteins are favourable candidates for immunotherapeutic targeting (*e.g.,* ERBB2-targeting ADCs for breast cancer (57)) and diagnostic imaging (*e.g.,* PSMA PET scans for prostate cancer (58)), we anticipate that our work can inform new avenues of translational research on HGSC. Expanding this surfaceome characterization to patient tumor- and ascites-derived cells will help reinforce the clinical relevance of these targets.

We focused on SLC7A1 for detailed characterization due to its strong and robust essentiality in multiple HGSC models. Interrogation of multi-omic patient datasets verified the cancer enriched expression profile of SLC7A1 observed in our *in vitro* models. We also demonstrate that increased SLC7A1 expression is associated with poorer overall survival in HGSC, suggesting that SLC7A1 may also have prognostic value. Our proteomic and functional characterization of SLC7A1 knockdown models uncovered critical dysregulations relating to cell growth and migration, protein synthesis and mitochondrial function culminating in caspase-mediated cell death in HGSC. While prior studies have reported associations between SLC7A1 and EOC, including immunohistochemical validation in patient samples and links to stromal activation, prior functional characterizations were conducted with models not reflective of the HGSC subtype and only included normal ovary surface epithelium cell controls (59,60). Provided that the different subtypes of EOC are well recognized as distinct disease entities (8), our careful utilization of models that recapitulate the unique molecular features of HGSC (16) augments the putative relevance of SLC7A1 in HGSC specifically. Further validation, including the development of high-affinity, high specificity reagents against SLC7A1 and evaluation of the heterogeneity of SLC7A1 essentiality and abundance in additional model systems and large patient cohorts, will be required to fully elucidate its prognostic and therapeutic utility in HGSC. SLC7A1 is one of four members of the cationic amino acid transporter (CAT) family and demonstrates the highest affinity for Arg (Km=0.1-0.16 mM) (46). In line with this, we show that SLC7A1 knockdown in HGSC results in reduced Arg transport into the cell and a decrease in mTORC1 signaling, a signaling cascade which requires Arg for activation. Analogous to SLC7A1 knockdown, depletion of Arg from cell culture media impaired HGSC proliferation suggesting that SLC7A1’s significance in HGSC is related to its arginine transporting function. Interestingly, we observed that SLC7A2, another member of the CAT family, albeit with lower affinity (Km=3.4-3.9 mM) and less specificity for Arg (43), similarly has increased abundance on the surface of HGSC cells compared to non-cancerous FTE cells and was critical for HGSC growth and survival, further implicating Arg transport as an important process in HGSC. Arg is an amino acid involved in several important biological processes such as protein translation, polyamine synthesis and mitochondrial function (52). Though Arg normally can be synthesized endogenously, it is well established that certain cancer types differentially produce and/or utilize Arg, thus necessitating increased transport of Arg from extracellular sources (52,61,62).

Correspondingly, we hypothesized that the increased surface abundance of Arg transporters in HGSC may similarly be reflective of dysregulated Arg metabolism. Indeed, our work revealed that HGSC and FTE cells had differential ability to utilize Arg precursor metabolites, specifically Cit, to sustain proliferation in the absence of Arg. We also uncovered that several enzymes involved in Arg metabolism are differentially abundant between HGSC and FTE cells in both *in vitro* models and patient samples. Most notably, ASS1 – the enzyme which metabolizes Cit – was found to be downregulated in HGSC cells, consistent with their inability to utilize Cit during Arg starvation in our study. Interestingly, depletion of ASS1 has been shown to be a deliberate strategy employed by other cancer types to divert cellular aspartate stores from arginine synthesis and instead fuel pyrimidine synthesis for increased proliferation (63). While our observation that SLC7A1 knockdown results in reduced DNA synthesis correlates with such a mechanism, further work will be required to clarify the underlying factors behind dysregulated Arg metabolism in HGSC. Additionally, multi-layered molecular profiling (*e.g*. surfaceome, phosphoproteome, metabolome *etc.)* following SLC7A1 depletion may reveal further biological insights into its role in HGSC. Overall, our work provides the first evidence for differential arginine metabolism between HGSC and FTE cells.

The discovery that certain cancer types rely on external sources of Arg has prompted the development of Arg deprivation therapies (64). Of note, enzymatic Arg depletion using formulations of recombinant ADI or arginase (*e.g.,* ADI-PEG20, PEG-rhARG1 and cohARG1-PEG) has demonstrated promising anti-cancer activity in early-stage clinical trials for mesothelioma and hepatocellular carcinoma (64). Equally significant, these trials have indicated minimal toxicity profiles underscoring the safe tolerance of therapeutically targeting Arg metabolism in patients (65–67). In our study, we show that ADI had deleterious effects on HGSC proliferation but no notable effects on non-cancerous FTE cells, providing evidence for targeting Arg metabolism in HGSC. Our study highlights Arg metabolism as an actionable liability in HGSC compared to its normal counterparts and warrants future investigation into therapeutically acting on this susceptibility in HGSC.

## CONCLUSIONS

In summary, we have performed a surface proteome characterization of HGSC using a panel of *in vitro* models and identified multiple HGSC enriched surface protein candidates. Our work underscores SLC7A1 as a putative clinically relevant surface protein and more broadly, sheds light on dysregulated Arg metabolism as a vulnerability in HGSC. As several other solute carrier proteins also demonstrated cancer enriched surface expression, we anticipate that our data may help reveal additional metabolic dysfunctions in HGSC. Beyond HGSC, the integrative framework employed in this study - coupling surfaceome profiling with systematic target prioritization and functional interrogation - could inform similar efforts to uncover novel surface protein dependencies in other cancer contexts.

## METHODS

### In vitro models

Cell lines were cultured as previously described (38). Briefly, commercial HGSC cell lines were provided by the Hakem lab and authenticated using Short Term Repeat DNA profiling (The Hospital for Sick Children, Toronto). Immortalized FTE cells were provided by the Rottapel lab and HGSC patient derived CAFs were isolated and provided by the Ailles lab. Cells were tested for mycoplasma contamination using the ATCC universal mycoplasma detection kit according to manufacturer’s instructions. Epithelial cells and CAFs were cultured in RPMI 1640 and IMDM, respectively, supplemented with 10% fetal bovine serum (FBS) and penicillin−streptomycin−glutamine (PSG) (100 U/mL penicillin, 100 μg/mL streptomycin, 292 μg/mL L-glutamine, Gibco). Epithelial cells and CAFs were incubated at 37 °C in 5% CO_2_/21% O_2_ and 2% O_2_, respectively.

### Proteomics sample preparation

#### Cell surface capture (CSC)

Cell lines were grown in 15 cm plates and CSC was performed when cells reached ∼90% confluency. As previously described (68,69), all cell surface labeling steps were performed on ice. Live, adherent cells were washed twice with washing buffer (PBS, 0.1% FBS), twice with labeling buffer (PBS, pH 6.5, 0.1% FBS) and subjected to oxidation with 1 mM sodium metaperiodate (NaIO_4_) in 20 mL of labeling buffer with slight rocking in the dark at 4°C for 15 min. Excess NaIO_4_ was removed with two washes of labeling buffer and cells were subsequently labeled with 5 mM biocytin hydrazide (Biotium, Fremont, CA, USA) in labeling buffer at 4°C with gentle rocking for 1 h. Cells were washed twice with wash buffer before being scraped off the plates and lysed in 1 mL of lysis buffer (40% 100 mM NH_4_HCO_3_, 40% 5X invitrosol, 20% acetonitrile) with 10 cycles of pulse sonication using an indirect ultrasonicator (Hielscher VialTweeter).

The BCA assay (Pierce) was used according to manufacturer’s recommendations to determine protein concentration and 2 mg of protein lysate was used for sample processing. Proteins were reduced with 5 mM dithiothreitol at 60°C for 30 min, alkylated with iodoacetamide in the dark for 30 min at room temperature (RT) and digested overnight for 16 h at 37°C with a Trypsin/Lys-C mix (Promega) at a 1:50 (w/w) enzyme:protein ratio. The next morning, digestion was quenched with 1X protease inhibitor cocktail (Roche). Peptides were combined with 100 µL of Streptavidin Sepharose™ High Performance beads (GE HealthCare) in 100 mM NH_4_HCO_3_, pH 8.0, and incubated for 1 h at RT with constant rotation. Beads were washed sequentially with 0.1X invitrosol, 5 M NaCl, 100 mM sodium carbonate (Na_2_CO_3_), pH 11, 80% isopropanol, and 100 mM NH_4_HCO_3_. *N*-glycopeptides were enzymatically deglycosylated and released from the beads with 5 U of PNGase F (Roche) in 100 µL of 100 mM NH_4_HCO_3_ at 37 °C for 16 h. Formerly *N*-glycosylated peptides were desalted using SP2 (70).

#### Whole cell lysate proteomics

Cells were washed 3X with cold PBS and scraped off cell culture plates in SP3 lysis buffer (100 mM HEPES, 1% SDS), heated for five min at 95 °C and underwent 10 cycles of pulse sonication. Protein concentration was quantified using a BCA assay (Thermo) according to manufacturer’s instructions and 50 µg of protein lysate was using for proteomic sample processing. Proteins were reduced with 5 mM dithiothreitol at 60°C for 30 min and alkylated with iodoacetamide in the dark for 30 min at RT. Proteins were digested using a magnetic bead based SP3 protocol (71). Briefly, Sera-Mag Speedbeads (Cytivia) were combined with protein lysates (10:1 [w/w] ratio) and protein binding was induced with ethanol at a final concentration of 70%. Supernatant was removed and magnetic beads were washed twice with 80% ethanol. The beads were then resuspended in 50 µL of 100 mM ammonium bicarbonate and Trypsin-LysC (Promega) was added at a 1:50 (w/w) enzyme:protein ratio for 16 h digestion at 37 °C. The following morning, the digestion was stopped with 0.1% trifluoroacetic acid. Peptides were desalted with C18 stage-tips and eluted peptides were dried in a centrifuge vacuum concentrator. Peptides were resuspended in 0.1% formic acid in LC-MS grade water and concentration was determined using a NanoDrop 2000 (Thermo Scientific).

### LC-MS/MS-based proteomics

LC-MS/MS data were acquired on an Orbitrap Q-Exactive (Thermo Fisher Scientific) coupled to an Easy-nLC 1000 nano-flow liquid chromatography system (Thermo Fisher Scientific). A two-column set up – with an Acclaim^TM^ PepMap^TM^ 100 column as the trap column and a 50 cm EasySpray^TM^ ES903 column (Thermo Fisher Scientific) for peptide separation – was used for liquid chromatography. Mass spectrometry was performed in positive-ion, data-dependent mode. Specific LC-MS/MS parameter details can be found in **Additional File: Table S4**.

### Proteomic data processing

#### CSC

MaxQuant (v1.6.3.3) (72) was used to search raw files against a UniProt human canonical plus isoform protein database (42,041 sequences) with the following search parameters enabled: match between runs, allowance of a maximum of two missed cleavages, carbamidomethylation of cysteine specified as a fixed modification and oxidation of methionine and deamidation of asparagine to aspartic acid (as a result of PNGase F elution) indicated as variable modifications. The false discovery of peptides was controlled using a target-decoy approach based on reversed sequences, and defined as 1% at site, peptide, and protein levels. CAF and epithelial samples were searched separately to minimize artificial matching between the two distinct cell types. The MaxQuant Asn-AspSites.txt output files were parsed into an in-house database for *N*-glycopeptide filtering and protein grouping. Peptides detected with an asparagine deamidation modification within the *N*-glycosylation *N*-[!*P*]-*STC* sequon (*N* = asparagine; [!*P*] = any amino acid other than proline; *STC* = serine, threonine, or cysteine at the +2 site) and with a localization probability > 0.8 were considered *N*-glycopeptides and were used in subsequent analyses. *N*-glycopeptide intensities were summed for *N*-glycoprotein quantification. Data were log_2_ transformed and median normalized and where necessary, missing protein intensities were imputed with random values between 1-1.2.

#### Global proteomics

MaxQuant (v1.6.3.3) (72) was used to search raw files against a UniProt human canonical plus isoform protein database (42,041 sequences) with the following search parameters enabled: match between runs, allowance of a maximum of two missed cleavages, carbamidomethylation of cysteine as a fixed modification and oxidation of methionine as a variable modification. The false discovery of peptides was controlled using a target-decoy approach based on reversed sequences, and defined as 1% at site, peptide, and protein levels. iBAQ values from the ProteinGroups.txt output file were used for analysis. Data were log_2_ transformed and median normalized.

### Bioinformatic analyses

Where appropriate, details regarding quantitative analyses (*e.g.,* statistical tests used, exact value of n) can be found in the relevant methods sections and figure legends. Significance levels can be found in the respective main text and figures. Statistical analysis and data visualization was performed in R (v.4.3.0) with the following packages: *ggplot2*, *ggpubr*, *reshape2*, *stringr*, *ggbeeswarm*, *vennDiagram*, *survival*, *survminer, lme4* and *lmerTest*.

#### Protein annotation

CIRFESS (14) was utilized to determine surface predictions (*i.e.* surface membrane and signal peptide predictions). Specifically, a *N*-glycoprotein was considered to have a predicted surface membrane localization if its surface prediction consensus (SPC) score was > 0 and a predicted signal peptide if it was predicted by at least one of three algorithms. *N*-glycoproteins were annotated as a plasma membrane protein if they were assigned a UniProt keyword of “Cell membrane” or “GPI-anchored” and a secreted protein if they were assigned the “Secreted” keyword. Surface protein family annotations were downloaded from HGNC and mapped by HGNC gene symbols.

#### Target prioritization

A two-staged approach was employed to prioritize HGSC enriched surface proteins. First, the CSC dataset was filtered for “high-confidence” HGSC surface proteins. This refers to proteins with a SPC score ≥ 1 (73), detected on the surface of ≥ 3 HGSC cell lines and detected in ≥ 1 HGSC tumors profiled by CPTAC (22,23). Secondly, a target scoring system was devised to rank the remaining 274 “high-confidence” proteins based on three criteria: 1) cancer enriched surface expression, 2) limited detection in normal tissue and 3) frequent detection in HGSC patient tissue. The first scoring criteria (*i.e.,* cancer enriched surface expression) leveraged our CSC data where targets were rewarded 1 point for each HGSC cell line that it had a log_2_FC > 1 compared to normal FTE cells and penalized 1 point for each HGSC cell line that it had a log_2_FC < -1 compared to normal FTE cells. Targets also received 1 point if it had an overall log_2_FC > 1 between HGSC cells and CAFs. Thus, enabling a maximum score of 5 for cancer enriched surface expression. For the second scoring criteria (*i.e.,* limited detection in normal tissue), we integrated four normal tissue protein datasets (9,24–26) to evaluate normal tissue detection. For each dataset, a protein was provided a score 0-1 indicating the proportion of normal tissue the protein was not detected in (*e.g.,* if a protein was not detected in 70% of normal tissue profiled in a single dataset, it would receive a score of 0.7). This entailed a maximum score of 4 across the four datasets. For the last scoring criteria (*i.e.,* frequent detection in HGSC patient tissue), a score ranging from 0-1 was assigned based on the proportion of HGSC tumors profiled by CPTAC (22,23) that a protein was detected in to ensure that prioritized targets were robustly detected in patient tissues (*e.g.* if a protein was detected in all 289 HGSC tumors profiled by CPTAC, it would receive a point of 1). In sum, the target score can range from 10 being the most cancer enriched surface protein and -4 being the least.

#### Multi-omic interrogation of SLC7A1 in patient datasets

Data utilized to investigate SLC7A1 protein abundance in bulk HGSC patient tumors and fallopian tubes were downloaded from supplementary tables 1 and 2 from McDermott *et al.,* 2020 (23). Data was used as is and no additional processing was performed.

Normalized read count and clinical annotation data pertaining to SLC7A1 transcript expression in HGSC patient tumors, precursor lesions and normal FT cells were acquired from supplementary tables 2 and 8 from Kader *et al.,* 2024 (39). For simplicity, regions annotated in the original study as “FT” or “fimbriae” were categorized as “Normal FT”, whereas “STIC” and “p53 signatures” regions were grouped as “Precursor lesions” and “Inv Cancer” and “Cancer (floating)” were considered “HGSC”. Data were log_2_ transformed for analysis.

Data used to characterize SLC7A1 protein abundance in EOC patient tumors of varying histological subtypes, benign ovarian masses and normal ovaries were obtained from supplementary tables 1 and 2 provided in Qian *et al.,* 2024 (42). Tumor (different samples from the same patient tumors) and LC-MS/MS injection (peptides from one sample run at different times) replicates were merged based on median values and were analyzed as a single patient. No additional data processing was performed.

To investigate associations between SLC7A1 transcript expression and overall survival, clinical patient and SLC7A1 RNA-seq expression data for the TCGA-OV cohort (43) was queried using FirebrowseR (v.1.1.35) (74). Log_2_ transformed RSEM quantification was used for analysis. SLC7A1 transcript expression was median dichotomized to define SLC7A1 high and low patients and a log-rank test was used to assess statistical significance for overall survival.

Kaplan-Meier curves were visualized using the packages *survival* (v.3.5.5) and *survminer* (v.0.4.9). Multivariate survival analysis was performed on the subset of patients that had clinical information recorded for all evaluated variables. Age was treated as a continuous variable and clinical stage as a categorical in reference to stage II.

To evaluate SLC7A1 mRNA expression in normal tissues, processed gene-level TPM data were downloaded from the online GTEx data portal (2017-06-05-V8) and data was used as is for visualization, with no additional processing.

### Knockdown pathway analysis

To evaluate proteomic consequences of SLC7A1 KD on OVCAR8 cells, GSEA (v.4.3.3) (75) was performed using a pre-ranked list of proteins sorted based on log_2_ fold change comparing cell treated with each SLC7A1 siRNA individually against cells treated with a scramble siRNA against Gene Ontology: Biological Processes. GSEA outputs were filtered for enriched gene sets that maintained consistent directionality across all three siRNAs (*i.e.,* |NES| > 1 in siA, siB and siC) and visualized in Cytoscape (v.3.9.1) using EnrichmentMap (v.3.3.6) and AutoAnnotate (v.1.4.0) modules.

### siRNA transfection

A reverse-transfection protocol was used for transient downregulation of targets. Three siRNA duplexes per gene and a scrambled siRNA duplex (negative control) were obtained from Origene (**Additional File: Table S4**). Lipofectamine RNAiMAX transfection reagent (Thermo) and siRNA duplexes were individually diluted with Opti-MEM transfection media (Gibco) and incubated for 5 min at RT. RNAiMAX was then gently combined with each siRNA mixture and incubated for 20 min at RT. The mixtures were then added to cell culture plates to ensure a final concentration of 10 nM (OVCAR8 and PEO4) or 5 nM (Kuramochi) of siRNA upon the addition of cells. Transfection reactions and cell densities were scaled based on the size of cell culture plate necessary for respective experiments.

### shRNA downregulation

Control (enhanced green fluorescent protein [GFP]) and SLC7A1 shRNAs were cloned into the pLKO-Tet-On lentiviral vector(76) (a gift from Dmitri Wiederschain, Addgene plasmid # 21915; http://n2t.net/addgene:21915; RRID:Addgene_21915), as per supplier’s recommendations. shRNA sequences were obtained from the MISSION RNAi database as follows:

SLC7A1 shRNA-1: 5’-CTGGGCTAATTGTGAACATTT-3’

SLC7A1 shRNA-2: 5’-GCTGAGGATGGACTGCTATTT-3’

GFP shRNA: 5’-TACAACAGCCACAACGTCTAT-3’

For lentivirus production, HEK293T cells were seeded in low antibiotic (0.1x Pen/Strep) DMEM media at a density of 1.0 x 10^6^ cells/well in 6-well plates. The next day, cells were co-transfected with 850 ng of packaging plasmid psPAX2 (psPAX2 was a gift from Didier Trono, Addgene plasmid # 12260; http://n2t.net/addgene:12260; RRID:Addgene_12260), 350 ng of envelope plasmid VSV-G(77) (pVSV-G was a gift from Akitsu Hotta, Addgene plasmid # 138479; http://n2t.net/addgene:138479; RRID:Addgene_138479), and 850 ng of pLKO-Tet-On shRNA constructs in OptiMEM using X-tremeGENE 9 DNA transfection reagent (Roche). The next day, media were replaced with viral harvest media (1% BSA, DMEM), followed by a final incubation for 24 h. Lentiviral supernatants were collected, passed through a 0.45 µm filter, and used immediately or stored at 4°C for a few days. For viral transduction 60,000 OVCAR8 cells/well were seeded in 6-well plates containing 8 µg/mL polybrene (Sigma). Following cell adherence, 100 µL of viral suspension was added to cells and incubated overnight. Media was replaced with puromycin containing media (3 µg/mL) for 48 h. Surviving, transduced cells were harvested for single cell cloning and characterization in subsequent experiments.

### Amino acid media modification experiments

RPMI 1640 medium for SILAC (Thermo) was prepared with 10% dialyzed FBS, PSG and L-lysine (final concentration of 0.22 mM as per the formulation for RPMI 1640). L-arginine was added in final concentrations relative to the amount of L-arginine regularly found in RPMI 1640 (*i.e.,* 100% Arg or complete RPMI = 1.15 mM Arg). Where applicable, citrulline (Cit) or argininosuccinic acid (AsA) was supplemented at a final concentration of 1.15 mM to match the amount of L-arginine normally found in RPMI 1640.

### Phenotype evaluations

#### Proliferation assay

3-5 replicates of 1000-2000 cells (depending on the cell line) were plated per experimental condition in 96-well plates. Cell growth was monitored using daily imaging with an Incucyte S5 Live-Cell Analysis System (Sartorius). 2D cell growth was quantified using the metric Phase Object Confluency to assess daily changes over a period of 7 days. Area under the curves (AUCs) for each growth curve were calculated and normalized against the mean negative control (either Scramble, GFP, complete RPMI media [CM] or vehicle control [VC]) AUC for the respective experiments. Statistical significance was determined by comparing AUCs of the respective conditions with a Student’s t-test.

#### Colony formation assay (CFA)

500-1000 cells per well were reverse-transfected with siRNA in 6-well plates, in triplicate. Media was changed after 24 h and plates were left undisturbed for 10-14 days following transfection. For shRNA experiments, dox (0.5ug/ml) was added to samples during seeding and media was replenished every 2-3 days. Cell densities and experiment duration were optimized for each cell line. At endpoint, plates were fixed and stained with 0.01% crystal violet, 20% methanol in PBS at RT for 2 h and washed. Plates were scanned using a Biotek Cytation Imaging Multimodal Reader (Agilent) and colonies were quantified using the Biotek Gen5 software (v3.11). Statistical significance was evaluated using a Student’s t-test between cells treated with each SLC7A1 siRNA against cells treated with a scramble siRNA.

#### Scratch wound migration assay

Following 24 h of siRNA transfection, OVCAR8 cells were detached, counted and re-seeded in Incucyte Imagelock 96-well plates (Sartorius) pre-coated with a thin-layer of rat collagen I at a density of 25,000 cells per well in triplicate. Cells were allowed to attach overnight. The next day, an Incucyte 96-Well Woundmaker Tool (Sartorius) was used to create reproducible wounds in each well. Wells were washed twice with PBS to remove dislodged cells, followed by the addition of media. Cell migration was monitored every 2 h following wound formation using the Scratch-Wound Analysis module on an Incucyte S5 Live-Cell Analysis System (Sartorius) and cell migration was quantified as percentage of the initial wound 10 h post wound. Statistical significance was evaluated using a Student’s t-test between cells treated with each SLC7A1 siRNA against cells treated with a scramble siRNA.

#### Spheroid growth assay

Following 24 h of siRNA transfection, OVCAR8 cells were detached, counted and re-seeded in round-bottom, ultra-low attachment 96 well plates (Corning) at a density of 5,000 cells per well in triplicate. Plates were centrifuged for 5 min at 300 *g* to pellet the cells at the bottom and placed in the incubator for 48 h to form spheroids. Once spheroid formation was visually confirmed, spheroids were embedded with 30 µL Matrigel (Corning). Spheroid growth was monitored every 48 h up to 10 days post transfection and media was changed as necessary. Spheroids were imaged using an EVOS inverted microscope (Life Technologies) and total area of spheroids was quantified using ImageJ (78). Statistical significance was evaluated using a Student’s t-test between cells treated with each SLC7A1 siRNA against cells treated with a scramble siRNA.

#### Annexin V staining

96 h post siRNA transfection, OVCAR8 cells were trypsinized, washed in PBS, counted and transferred to round bottom flow cytometry tubes at a density of 200,000 cells per tube. Three transfection replicates per condition were used for staining and flow cytometry. Cells were stained with an Annexin V Apoptosis Detection Kit (eBioscience) according to manufacturer’s recommendations. Cells were analyzed by flow cytometry using a BD FACSymphony^TM^ A3 Cell Analyzer and data were analyzed using FlowJo (v10.10). Statistical significance was evaluated using a Student’s t-test between cells treated with each SLC7A1 siRNA against cells treated with a scramble siRNA.

#### Puromycin incorporation

48 h post siRNA transfection, cell culture media was replaced with media containing 10 µg/mL puromycin and incubated for 30 min at 37 °C. OVCAR8 cells were trypsinized, washed in PBS, counted and transferred to round bottom flow cytometry tubes at a density of 200,000 cells per tube. Staining and flow cytometry was performed on three transfection replicates per condition. Cells were fixed and permeabilized using BD Cytofix/Cytoperm and BD PermWash, respectively. Cells were stained with an anti-puromycin antibody (clone 12D10 [Millipore Sigma], 1:200) for 1 h at RT, washed and then stained with an anti-mouse Alexa Fluor 488-conjugated secondary antibody (Cat# ab150113, Abcam) for 20 min at RT. Cells were analyzed by flow cytometry using a BD LSRFortessa^TM^ X-20 Cell Analyzer and data were analyzed as previously described (79) using FlowJo (v10.10). Statistical significance was evaluated using a Student’s t-test between cells treated with each SLC7A1 siRNA against cells treated with a scramble siRNA.

#### EdU incorporation

DNA synthesis was measured using the Click-IT^TM^ Plus EdU Flow Cytometry Assay Kit (Thermo Fisher) as recommended by the manufacturer with the following specifications. 72 h post transfection, cells were incubated with 10 µM EdU for 90 min and 500,000 cells were used per replicate. Three transfection replicates per condition were used for staining and flow cytometry. Cells were analyzed by flow cytometry using a BD LSRFortessa^TM^ X-20 Cell Analyzer and data were analyzed using FlowJo (v10.10). Statistical significance was evaluated using a Student’s t-test between cells treated with each SLC7A1 siRNA against cell treated with a scramble siRNA.

#### Mitochondria characterization

72 h post siRNA transfection, OVCAR8 cells were trypsinized, washed in PBS, counted and transferred to round bottom flow cytometry tubes at a density of 250,000 cells in 500 µL per tube. Cells were incubated with 0.6 µM TMRE (Thermo Fisher; Cat# T669) and 50 µM Verapamil (Tocris) or 15 µM MitoSOX Red (Invitrogen) for 30 min at 37°C. Flow cytometry was performed using the BD FACSCelesta^TM^ Cell Analyzer and data were analyzed using FlowJo (v10.10). Dead cells were excluded using DAPI following doublet exclusion. Three transfection replicates per condition were used for staining and flow cytometry. Statistical significance was evaluated using a Student’s t-test between cells treated with each SLC7A1 siRNA against cell treated with a scramble siRNA.

### *In vivo* xenograft growth

Young adult (6-8-weeks old) female immunodeficient NOD.Cg-Prkd^cscid^ Il2rg^tm1Wjl^/SzJ (NSG, Strain # 005557) mice from the Jackson Laboratory were used for experiments. Mice were housed in a modified barrier, specific pathogen-free facility in sealed negative ventilation cages (Allentown) in groups of maximum five mice per cage, at 22°C–24°C and a 12-h light/12-h dark cycle with food and water *ad libitum*. Animal experiments were performed according to guidelines from the Canadian Council for Animal Care and under protocols approved by the Animal Care Committee of the Princess Margaret Cancer Centre (Protocol number 6396). 1 x 10^6^ OVCAR8 cells were resuspended in 100 μl of equal volume growth factor reduced Matrigel (Corning) and RPMI media and injected subcutaneously into the right flank of each animal. Ten animals were injected per condition (two SLC7A1 shRNA containing clonal populations and one GFP shRNA containing clonal population). 10 days following engraftment, five mice per condition were administered drinking water containing doxycycline (1mg/ml) and 5% sucrose until endpoint (Dox group). The remaining five mice per control (non-treated [NT] group) were administered drinking water with 5% sucrose until endpoint. Tumors were measured biweekly using calipers and tumor volume (*V*) was calculated with the formula: *V* = 0.5 x *l* x *w*^2^, where *l* and *w* are the longest and shortest perpendicular measurements, respectively. As recommended by the ACC, endpoint was defined as when the non-treated mice in any condition reached approximately 15 mm in length. At endpoint, animals were sacrificed with CO_2_ and tumors were removed, measured, and weighed. AUCs for each growth curve were calculated and normalized against the mean NT for each respective condition. Statistical significance was determined by comparing relative AUCs between the respective conditions with a Student’s t-test.

### Quantitative PCR (qPCR)

RNA was isolated from cells using a RNeasy Mini Kit (QIAGEN) and reverse transcribed into cDNA using SuperScript^TM^ IV VILO Master Mix (Invitrogen) by following manufacturer’s recommendations. qPCR was performed using 10ng of cDNA, LUNA universal qPCR Master Mix and the following primers:

SLC7A1 forward: 5’ GCCTGTGCTATGGCGAGTTT 3’

SLC7A1 reverse: 5’ ACGCTTGAAGTACCGATGATGTA 3’

GAPDH: forward: 5’ ATGGCCTTCCGTGTCCCCACTG 3’

GAPDH: reverse 5’ GTGGGTGTCGCTGTTGAAGTCAG 3’

The QuantStudio 6 Pro (Thermo) was used for qPCR and analysis.

### Immunoblotting

Cell pellets were lysed in cold RIPA buffer (50 mM Tris-HCl pH 8, 150 mM NaCl, 5 mM EDTA, 1% NP-40, 0.1% SDS), supplemented with Protease Inhibitor Cocktail tablets (Roche) and Phosphatase Inhibitor Cocktail Mini tablets (Roche). 10-20 µg of lysate per well were resolved in 8-15% SDS-PAGE gels and wet-transferred to polyvinylidene fluoride membranes overnight. The next day, membranes were blocked for 1 h in 5% milk (w/v) in Tris-buffered saline Tween-20 and subjected to overnight incubation with primary antibodies (**Additional File: Table S4**) at 4 °C. Membranes were washed and incubated with HRP-conjugated secondary antibodies for 1 h at RT and immunorective bands were visualized with SuperSignal West Femto Maximum Sensitivity Substrate (Pierce) and a MicroChemi imager (Bio-Imaging Systems)

### LC-MS/MS based metabolomics

48h post siRNA treatment, OVCAR8 cells were washed once with PBS and incubated with RPMI media prepared with 1.15 mM ^13^C_6_ L-Arg for 1 minute at 37 °C. Following pulse labelling, cells were washed with ice-cold 150 mM ammonium formate, pH 7.4. Cells were then scraped and extracted with 600 μL of 31.6% MeOH/36.3% ACN in H_2_O (v/v). Cells were lysed and homogenized via sonication using a Bioruptor Pico (Diagenode) at 4 °C. Cellular extracts were partitioned into aqueous and organic layers following dimethyl chloride treatment and centrifugation. The aqueous supernatants were dried by vacuum centrifugation with sample temperature maintained at -4°C (Labconco, Kansas City MO, USA). Dried extracts were subsequently re-suspended in 50 μL of chilled H_2_O and clarified by centrifugation at 1°C. Sample injection volumes for analyses were 5 μL per injection. Four transfection replicates per condition were used for analysis.

For stable isotope tracer metabolite analysis, samples were analyzed using an Agilent 6545 quadrupole time of flight (QTOF)–LC-MS/MS. Chromatography was achieved using a 1290 Infinity ultra-performance LC system (Agilent Technologies, Santa Clara, CA, USA). Mass spectrometer was equipped with a Jet Stream electrospray ionization (ESI) source, and samples were analyzed in positive mode. Source gas temperature and flow were set at 325 °C and 9 L/min respectively, nebulizer pressure was set at 45 psi and capillary voltage was set at 4000 V. The autosampler temperature was maintained at 4°C.

Chromatographic separation of metabolites was achieved using an Intrada Amino Acid column 3 μm, 3.0×150 mm2 (Imtakt Corp, JAPAN). The chromatographic gradient started at 100% mobile phase B (0.3% formic acid in ACN) with a 3 min gradient to 27% mobile phase A (100 mM ammonium formate in 20% ACN / 80% water) followed with a 19.5 min gradient to 100% A at a flow rate of 0.6 mL/min. This was followed by a 5.5 min hold time at 100% mobile phase A and a subsequent re-equilibration time (7 min.) at initial conditions before the next injection. The column temperature was maintained at 10 °C.

Retention times and linear range of detection were obtained by running authentic standard mixes. Repeat injections of authentic standards were performed throughout the queue to observe any shifts in retention time or chromatographic quality. Area under the curve for each sample and metabolite and isotopes were analyzed and ensured to be below the saturation limit for those metabolites where range curves were available. No corrections or allowances were made for ion suppression effects. Stable isotope tracer data were analyzed, and natural abundance matrix correction performed using Profinder software (Agilent Technologies). Cells exposed to ^13^C-labeled nutrients as well as control (*i.e.,* unlabelled) OVCAR8 samples exposed natural abundance nutrients were used to ensure that observed ^13^C-incorporation into other metabolites is not an interfering ion.

## DECLARATIONS

### Ethics approval and consent to participate

Animal experiments were performed according to guidelines from the Canadian Council for Animal Care and under protocols approved by the Animal Care Committee of the Princess Margaret Cancer Centre (Protocol number 6396).

### Consent for publication

NA

### Availability of data and materials

All proteomics mass spectrometry raw files acquired in this study are available from UCSD’s MassIVE database under the dataset identifier #MSV000099007 (ftp://MSV000099007@massive-ftp.ucsd.edu, reviewer password: HGSCproteomics34). Data will be made public after the acceptance of the manuscript. Processed proteomics data are available in this paper’s **Tables S1 and S3**. Images of full immunoblot membranes can be found in **Figure S5**. Any additional information required to reanalyze the data reported in this paper is available from the corresponding author upon request.

### Authors’ contributions

M.G., M.W., S.M-G., B.L., T.L., and X.L. performed experiments. M.G. and V.I. performed data analysis. M.G. completed data visualization. T.K., L.A., C.L.J and A.N.T. supervised the research and provided resources. M.G. and T.K. wrote the manuscript and all authors contributed to editing and approved the final manuscript.

## Acknowledgements

The authors thank all members of the Kislinger lab for helpful suggestions and Dr. Hakem and Dr. Rottapel (Princess Margaret Cancer Centre) for providing epithelial cell lines. The Genotype-Tissue Expression (GTEx) Project was supported by the Common Fund of the Office of the Director of the National Institutes of Health, and by NCI, NHGRI, NHLBI, NIDA, NIMH, and NINDS. The results here are in whole or part based upon data generated by the TCGA Research Network. Metabolite measurements were performed at the Rosalind and Morris Goodman Cancer Research Centre Metabolomics Core Facility supported by the Canada Foundation for Innovation, The Dr. John R. and Clara M. Fraser Memorial Trust, the Terry Fox Foundation (TFF Oncometabolism Team Grant 116128), and McGill University.

## ADDITIONAL FILES

**Figures S1-S5**

**Table S1**: Processed CSC data.

**Table S2**: Target scoring of 274 HGSC enriched surface proteins.

**Table S3**: SLC7A1 knockdown OVCAR8 WCL proteomics

**Table S4**: Experimental details

---

**Figure S1:**
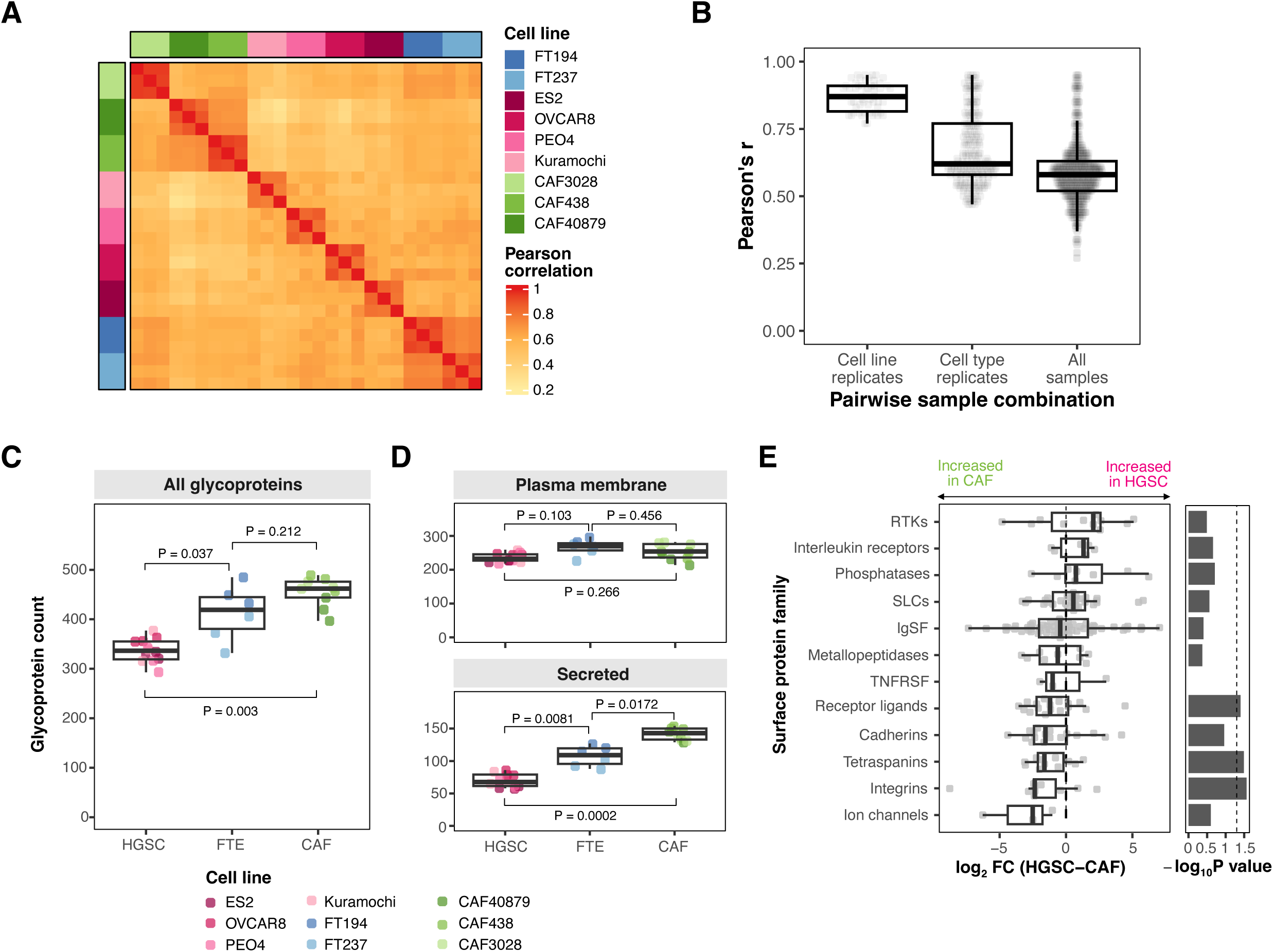
Surface proteomic characterization of HGSC.

**Figure S2:**
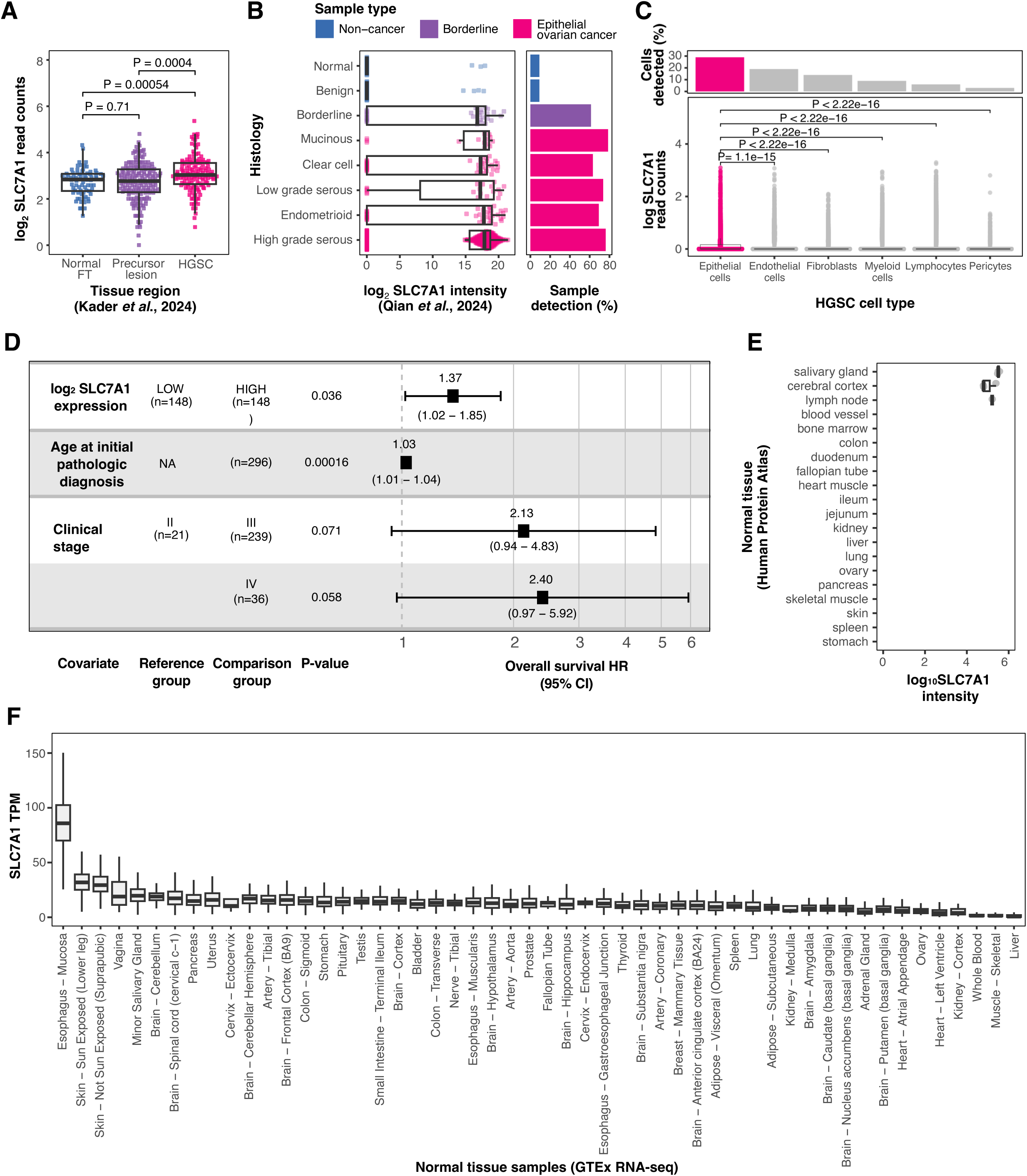
SLC7A1 is a cancer enriched surface protein.

**Figure S3:**
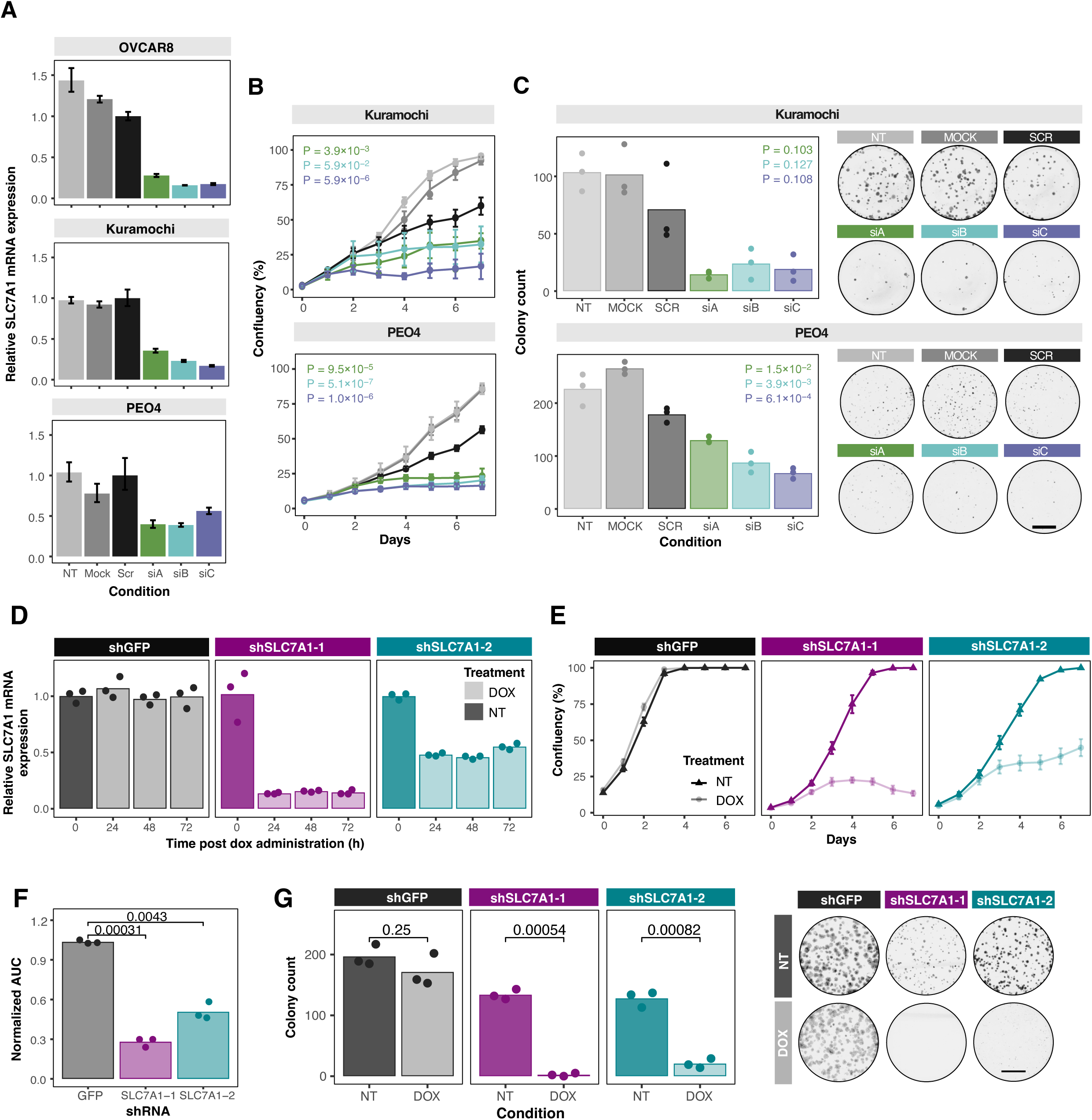
SLC7A1 is essential for HGSC growth.

**Figure S4:**
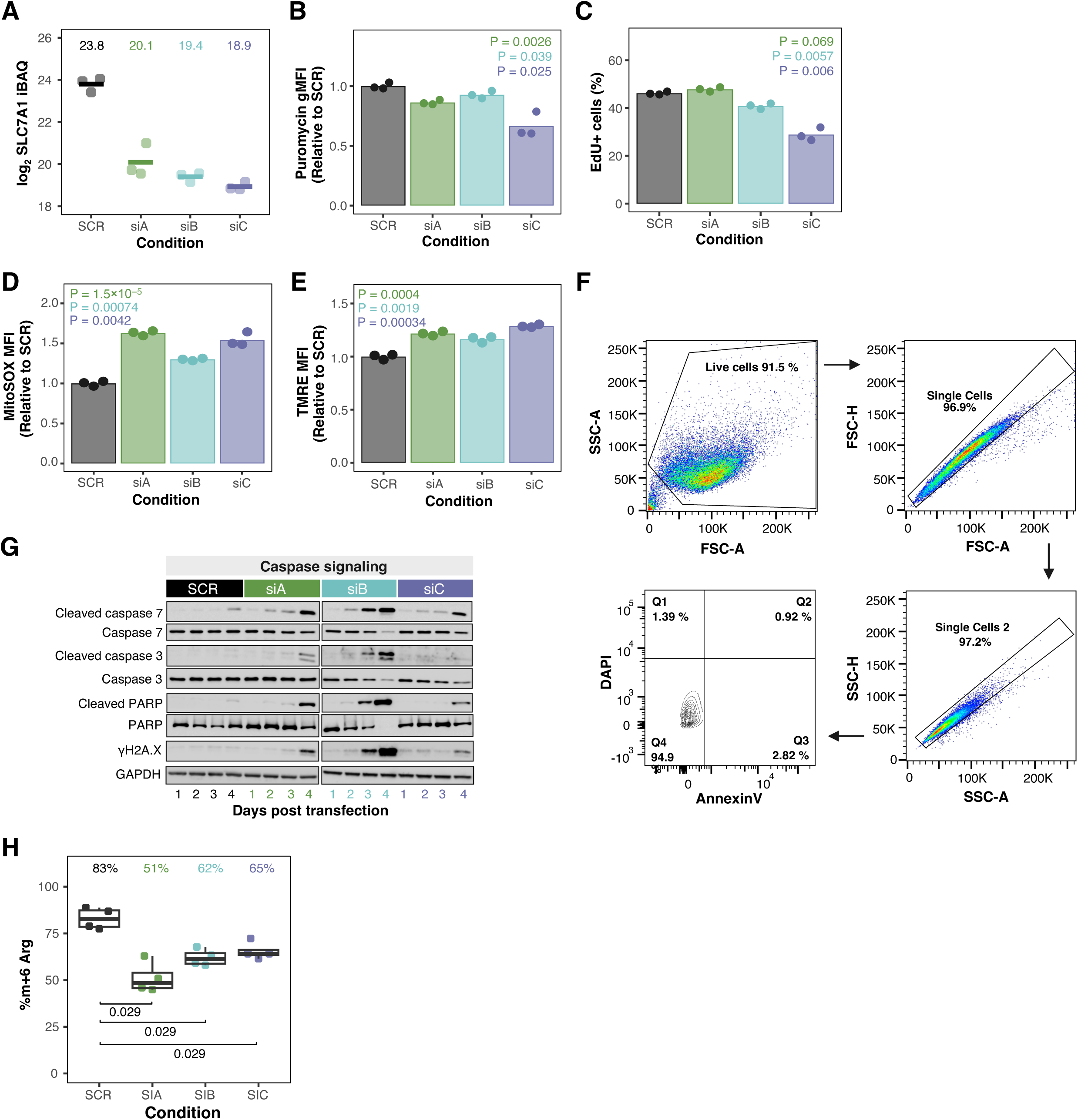
SLC7A1 knockdown impairs several cellular processes in HGSC.

**Figure S1:**
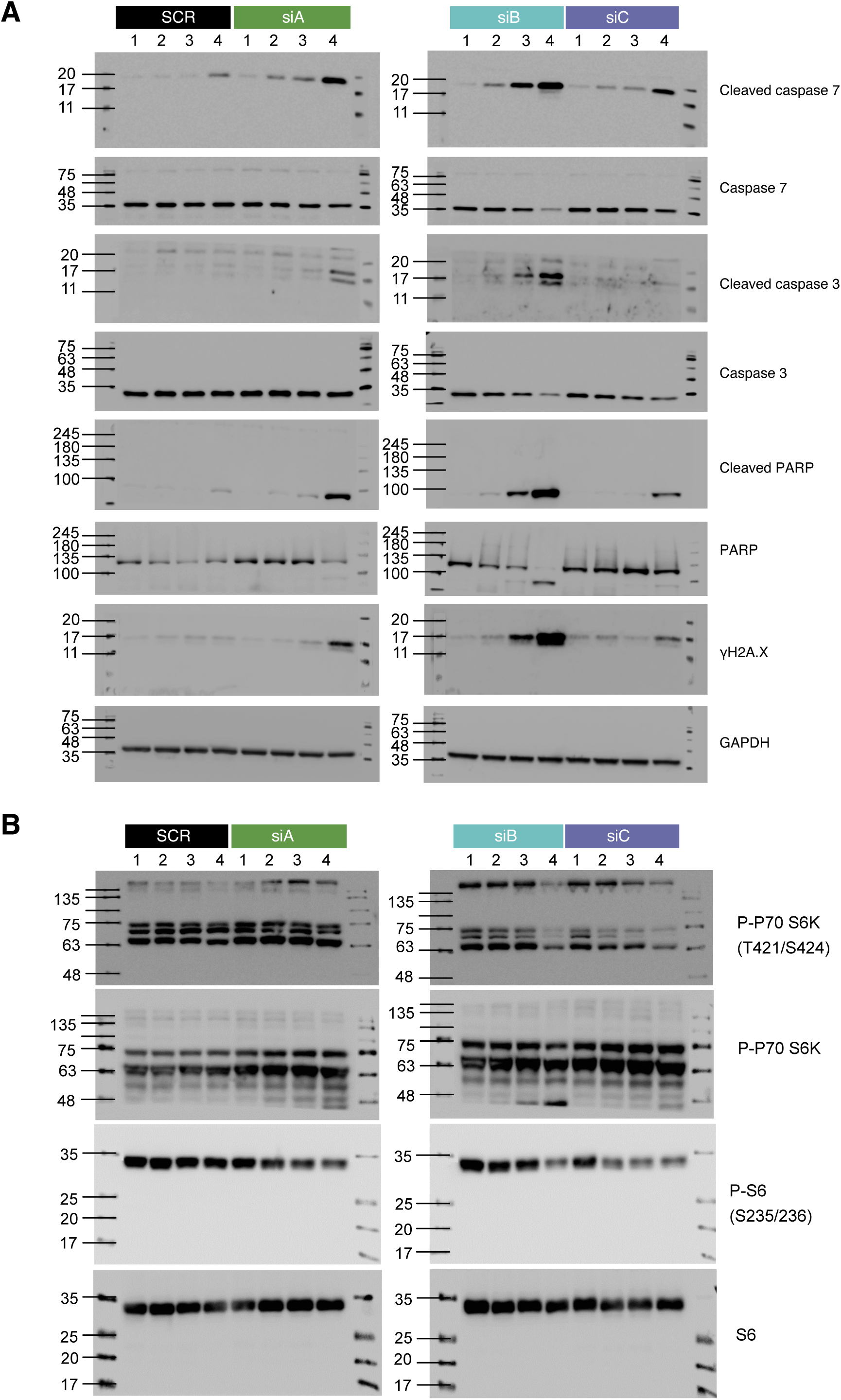
Full immunoblots.

