## Supplemental Figures for "Surface proteomics reveals arginine metabolism as a vulnerability in high grade serous ovarian cancer"

### Figure S1: Surface proteomic characterization of HGSC

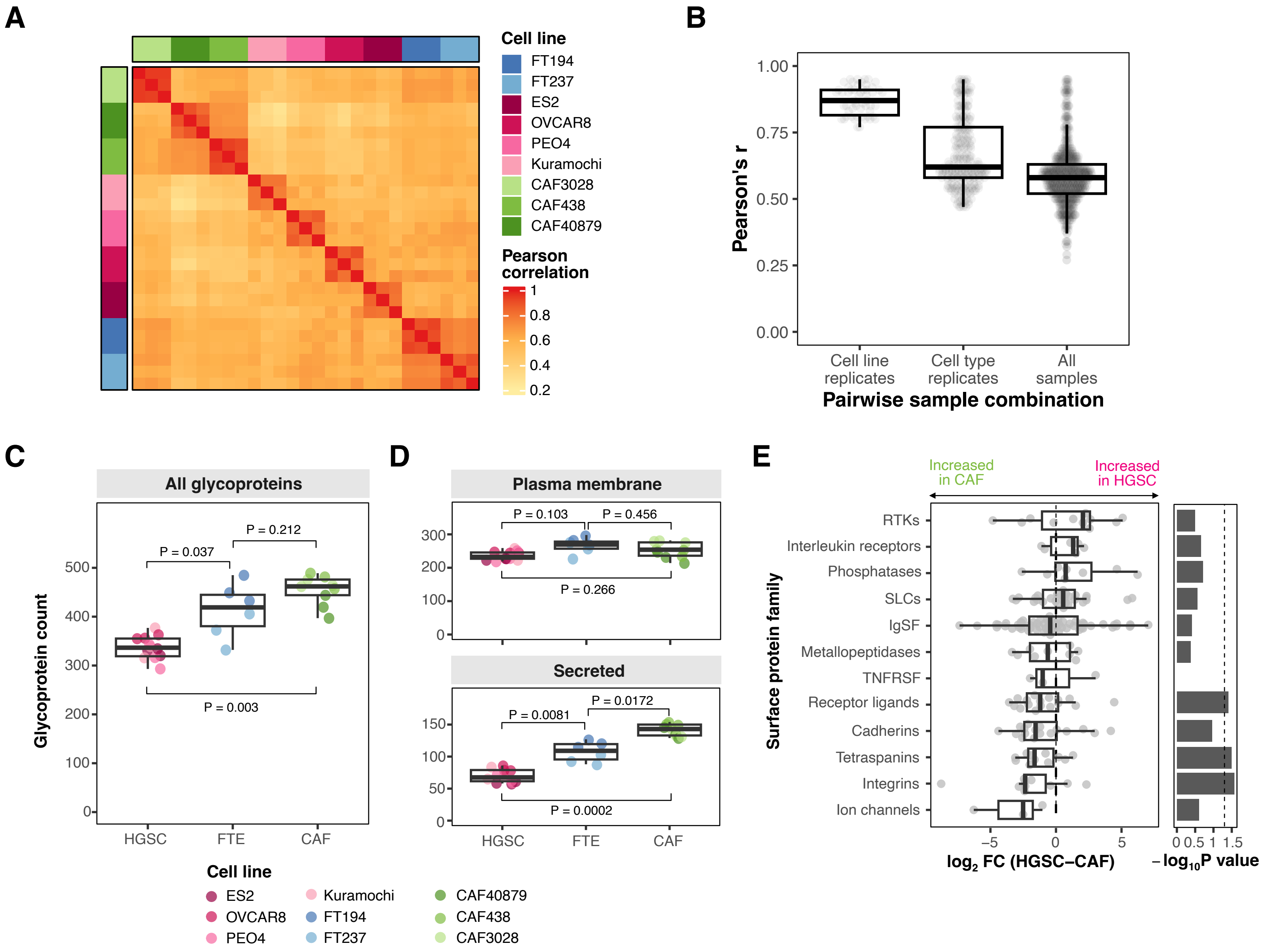

### Figure S2: SLC7A1 is a cancer enriched surface protein

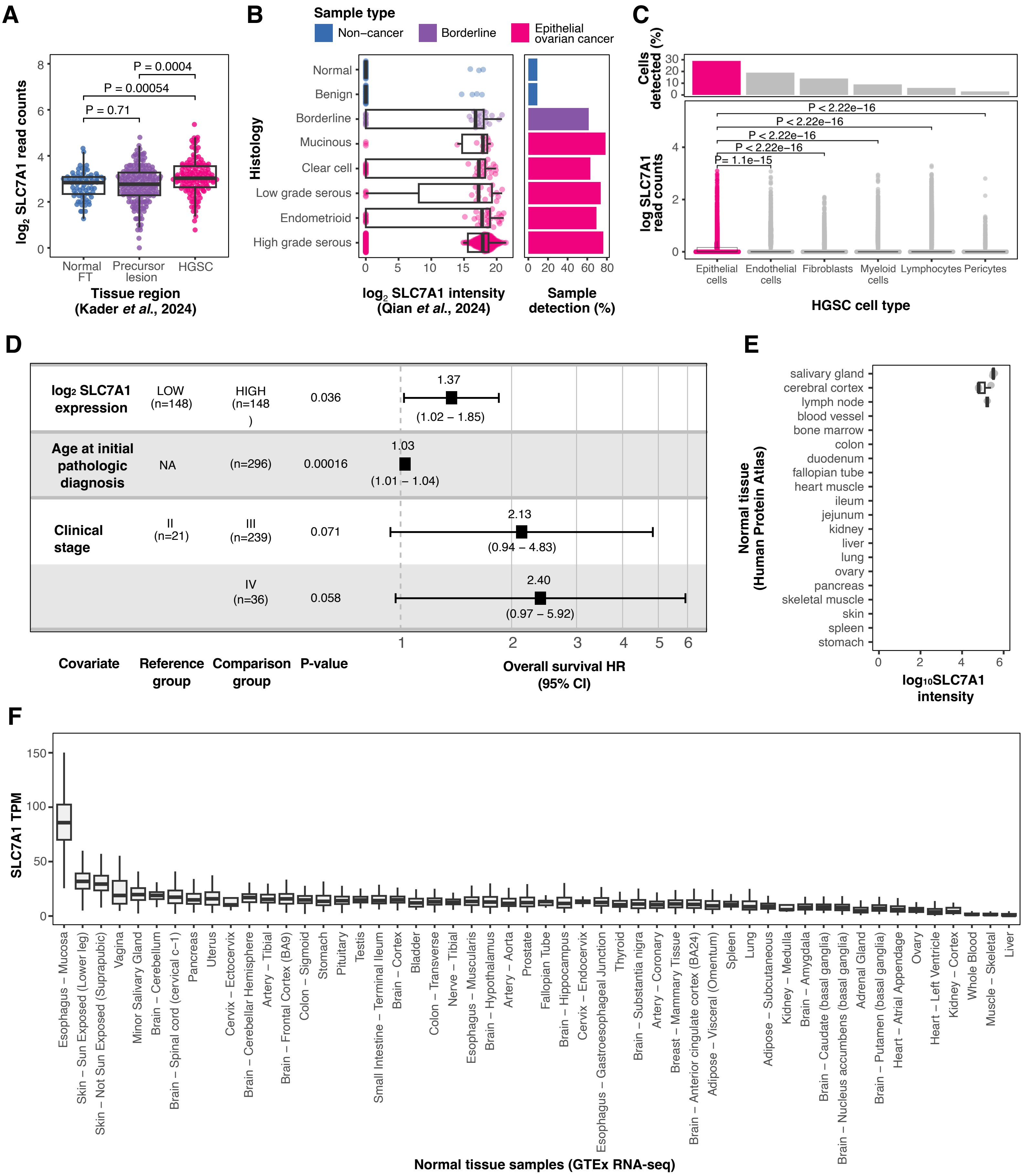

### Figure S3: SLC7A1 is essential for HGSC growth

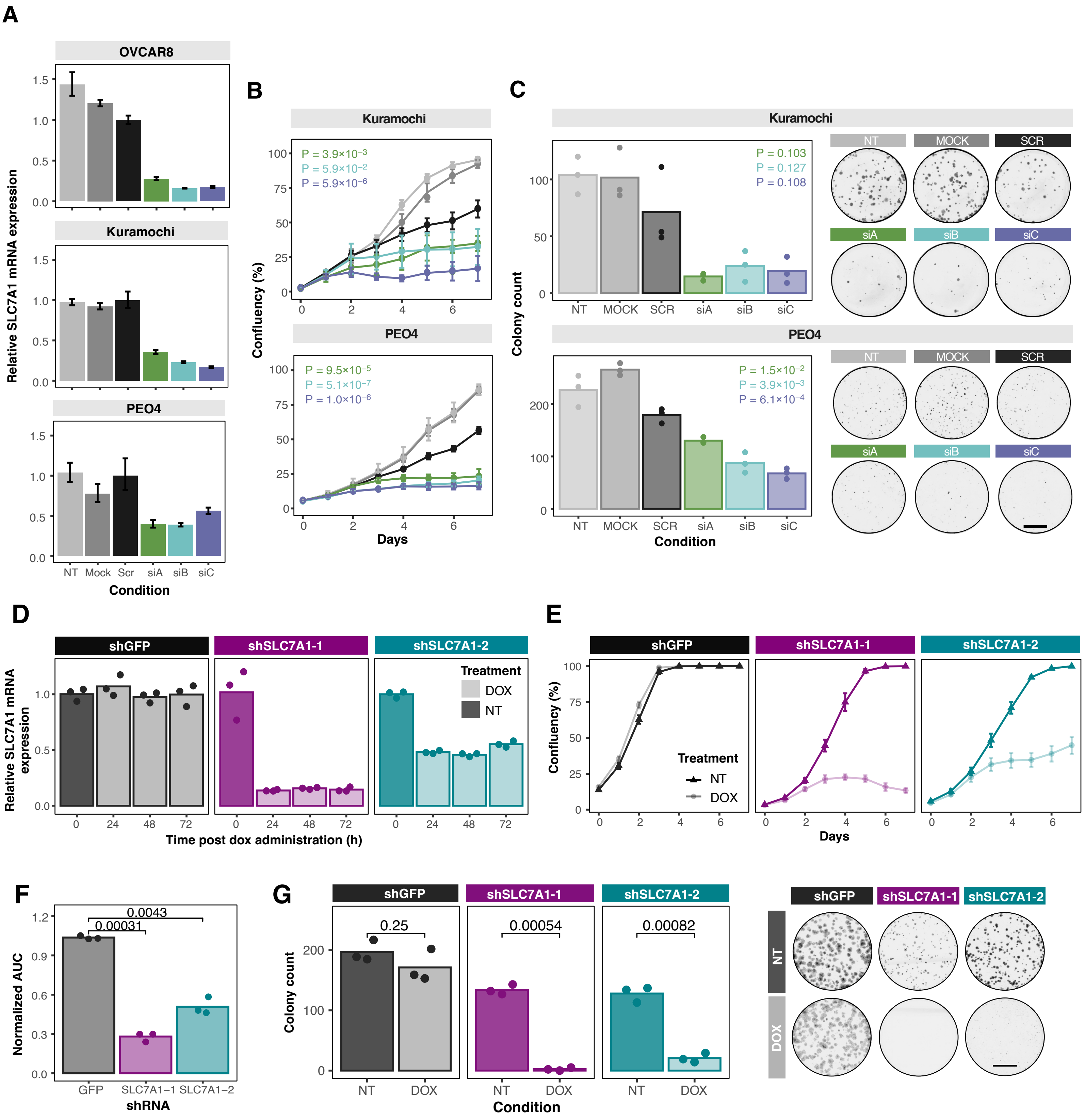

### Figure S4: SLC7A1 knockdown impairs several cellular processes in HGSC

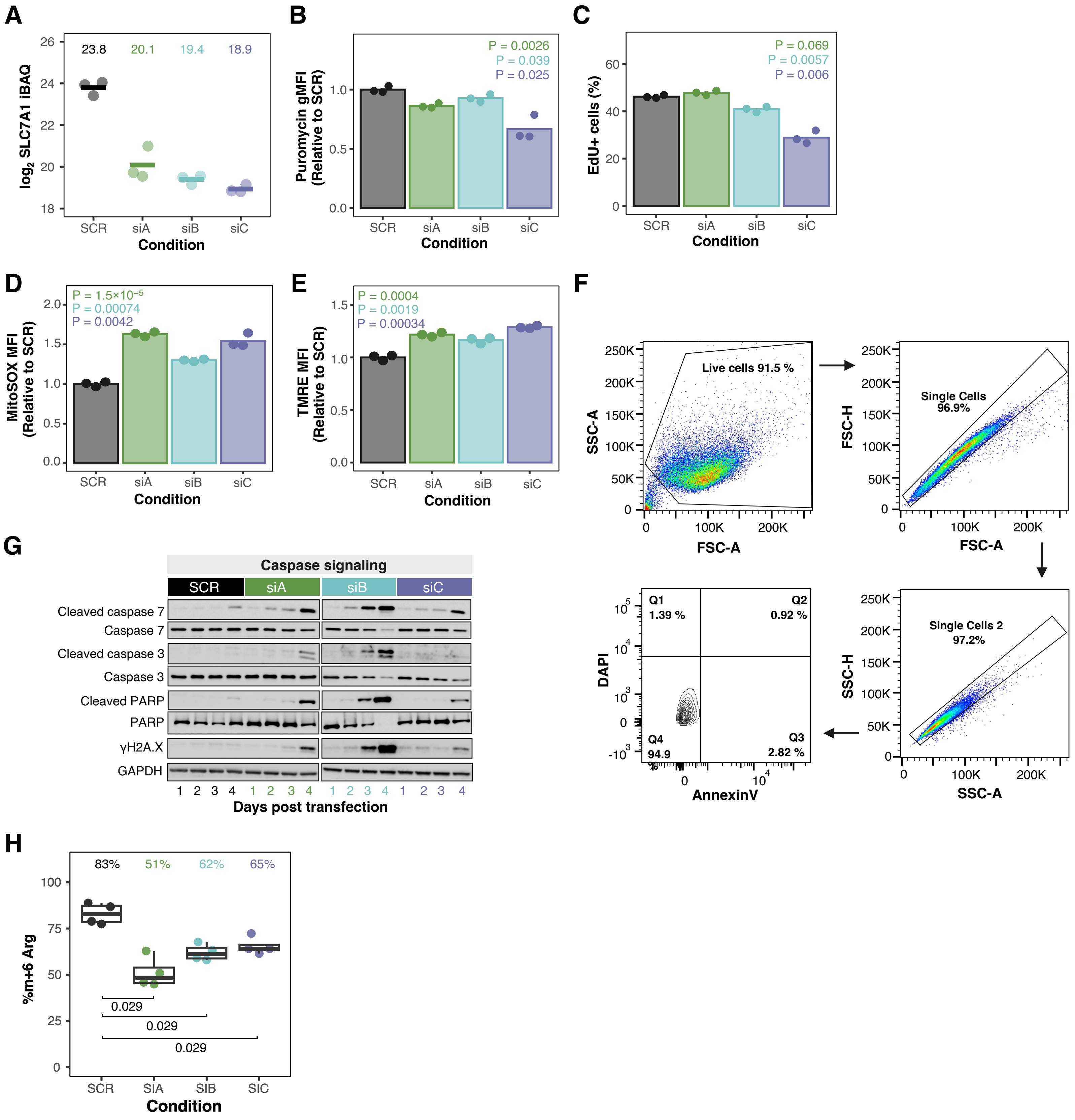

### Figure S5: Full immunoblots

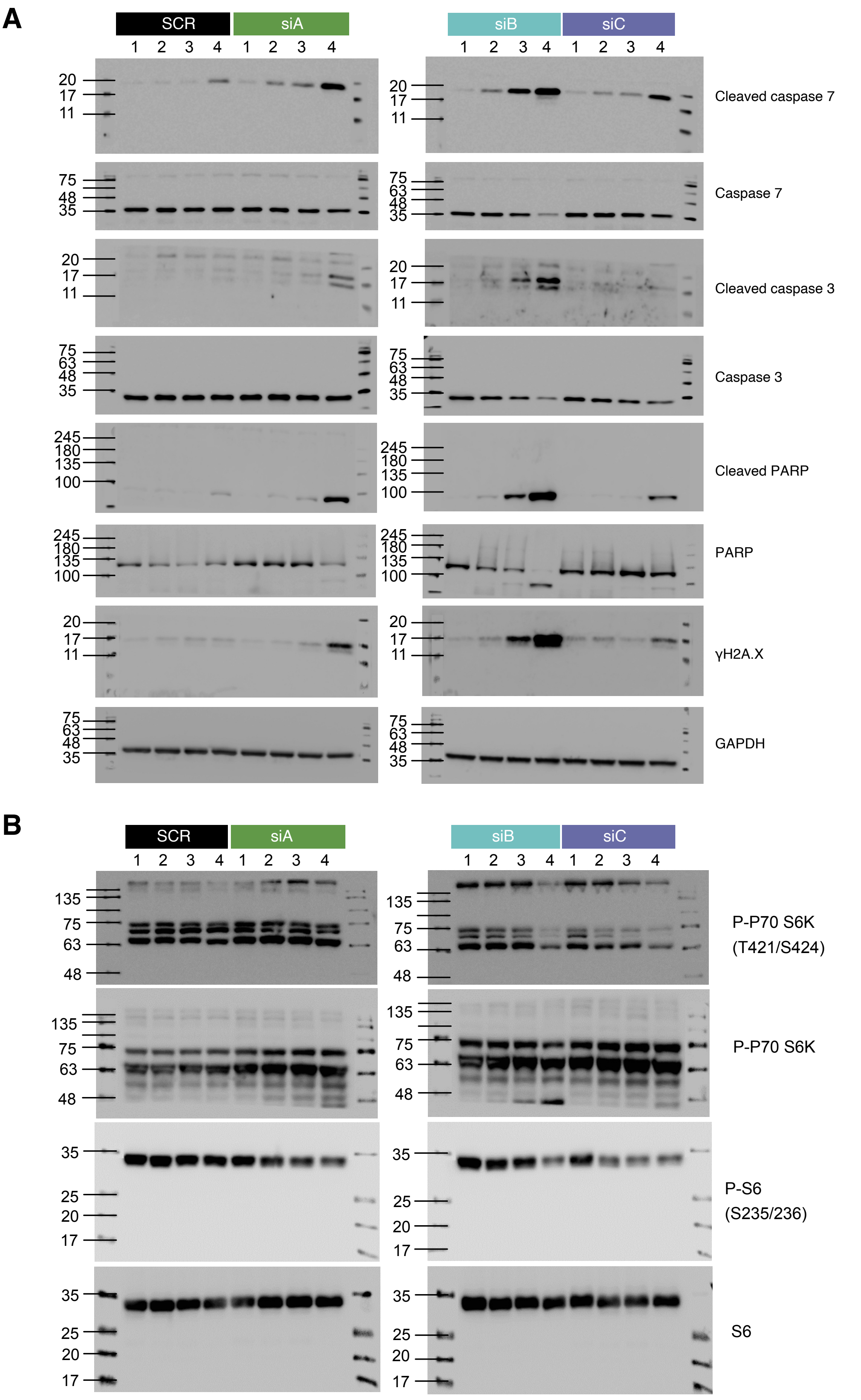
